# *In Situ* Surface-Directed Assembly of 2D Metal Nanoplatelets for Drug-Free Treatment of Antibiotic-Resistant Bacteria

**DOI:** 10.1101/2021.09.28.462217

**Authors:** Parinaz Fathi, Ayman Roslend, Maha Alafeef, Mandy B. Esch, Dipanjan Pan

## Abstract

The development of antibiotic resistance among bacterial strains is a major global public health concern. To address this, drug-free antibacterial approaches are needed. High-touch surfaces in particular can serve as a means for the spread of bacteria and other pathogens from one infected person to another. Copper surfaces have long been known for their antibacterial properties. To further enhance the surface’s antibacterial properties, we used a one-step surface modification technique to assemble 2D copper chloride nanoplatelets directly onto copper surfaces such as copper tape, transmission electron microscopy (TEM) grids, electrodes, and granules. The nanoplatelets were formed using copper ions from the copper surfaces, enabling their direct assembly onto these surfaces in a one-step process that does not require separate nanoparticle synthesis. The synthesis of the nanoplatelets was confirmed with TEM, scanning electron microscopy, energy dispersive spectroscopy (EDS), x-ray diffraction (XRD), and Fourier transform infrared spectroscopy (FT-IR). Antibacterial properties of the surfaces with copper chloride nanoplatelets were demonstrated in multi-drug-resistant (MDR) *E. coli*. The presence of copper chloride nanoplatelets on the surface led to a marked improvement in antibacterial properties compared to the untreated copper surfaces. Surfaces with copper chloride nanoplatelets affected bacterial cell morphology, prevented bacterial cell division, reduced their viability, damaged bacterial deoxyribonucleic acid (DNA), and altered protein expression. In particular, proteins corresponding to cell division, DNA division, and mediation of copper toxicity were down-regulated. This work presents a robust method to directly assemble copper chloride nanoplatelets onto any copper surface to imbue it with improved antibacterial properties. To demonstrate that our method of particle generation can be used with other metal surfaces, we also demonstrate the synthesis of other metal-derived nanoarchitectures on a variety of metal surfaces.

## INTRODUCTION

Drug-resistant bacterial infections are a rising concern with increasing prescription of antibiotics. High-touch surfaces have been proven to contribute to pathogen spread.^1^ Attempts to provide surfaces with antibacterial properties often rely on the application of surface coatings.^2^ Nanomaterials have been successfully synthesized for use in a variety of applications in medicine,^3–14^ including the detection and eradication of bacteria.^15–22^ Nanoparticle-based approaches to developing antibacterial surfaces typically rely on the synthesis of nanoparticles followed by their subsequent incorporation into polymer-based coatings that do not have inherent antibacterial properties.

The antibacterial properties of copper surfaces have long been known, and copper is used in a variety of medical devices including dental implants, intrauterine devices, and catheters.^23–28^ The goal of this work was to further improve the antibacterial properties of copper surfaces by developing antibacterial coatings that could be assembled directly onto copper surfaces. Using a robust and simple synthesis process, copper chloride nanoplatelets were assembled onto a variety of copper surfaces **(Figure 1)**. The synthesis was conducted by dropping a solution of dilute HCl or dilute HCl in combination with 2 2′-(ethylenedioxy)bis(ethylamine) directly onto the metal surface. The resulting nanoplatelets are referred to as Cu@HCl NP and Cu@HCl-NH_2_ NP, respectively. The physicochemical properties of the nanoplatelets were extensively characterized, and the improved antibacterial properties of the particle-covered surfaces were compared to those of untreated copper surfaces using data obtained with *E. coli* and Multi-drug-resistant (MDR) *E. coli*. These model bacteria were selected because drug resistance in *E. coli* is of increasing concern.^29–33^

**Figure 1.**
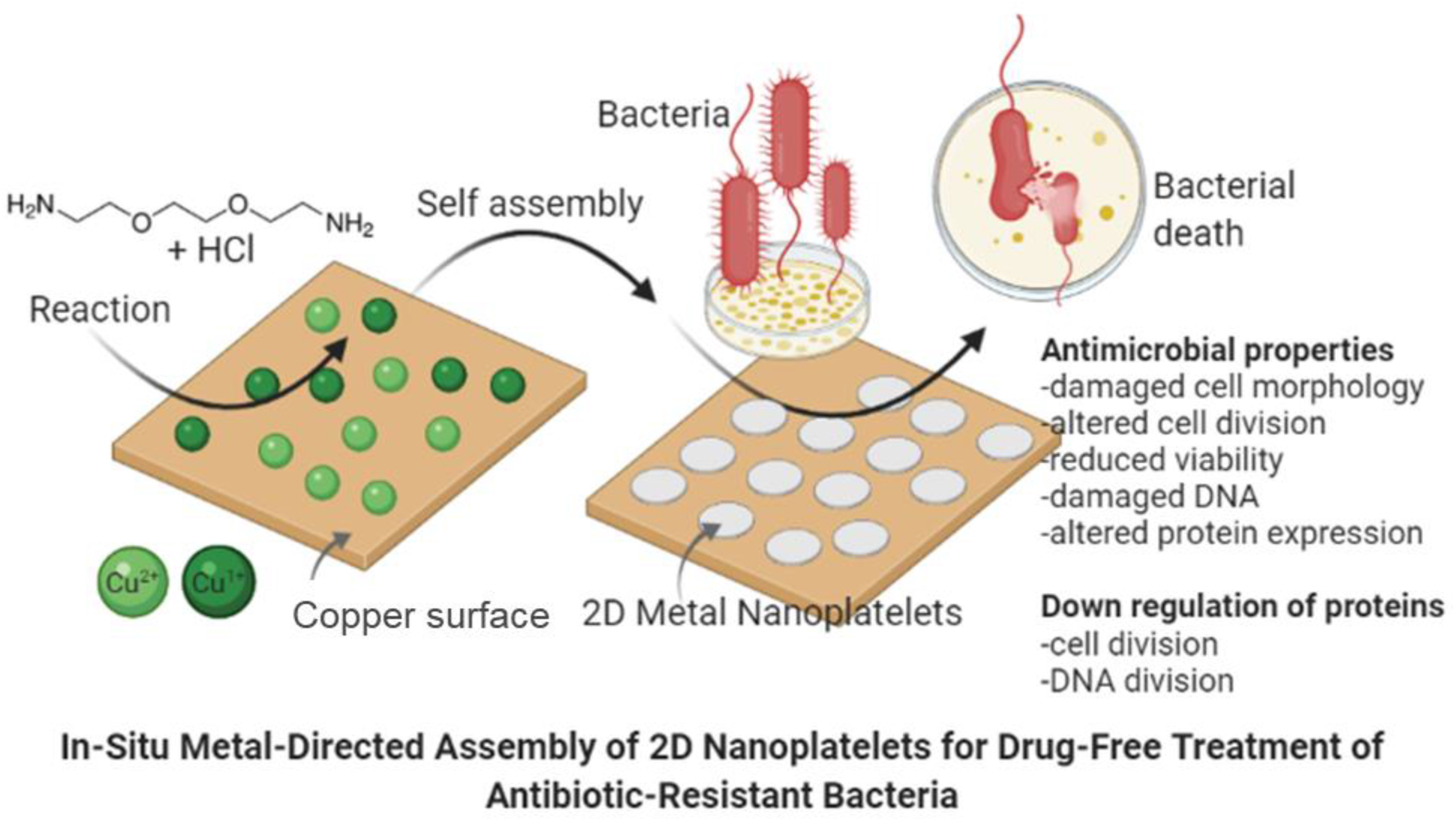
Self-assembly of copper-derived nanoplatelets onto copper surfaces. A synthesis solution composed of dilute HCl or dilute HCl in combination with small amounts of 2 2′-(ethylenedioxy)bis(ethylamine) is deposited onto a copper surface and allowed to dry. No copper is present in the synthesis solution itself. Upon deposition of the synthesis solution onto the copper surface, chlorine ions from the HCl couple with copper ions from the surface to form copper chloride crystals (copper nanoplatelets). The nanoplatelets exhibit improved antibacterial properties compared to the copper surfaces alone. The inclusion of 2 2′-(ethylenedioxy)bis(ethylamine) modifies nanoplatelet morphology.

## RESULTS AND DISCUSSION

### Copper nanoplatelets were assembled directly on multiple copper surfaces

The precursor solution for Cu@HCl NPs contained only dilute HCl, while the precursor solution for Cu@HCl-NH_2_ NPs contained dilute HCl and a small amount of 2 2′-(ethylenedioxy)bis(ethylamine) (referred to as diamine). Full details of experimental procedures are provided in the Methods section. Deposition of the solutions onto the copper side of copper TEM grids led to the formation of nanoplatelets directly on the TEM grid surfaces. TEM imaging revealed the formation of Cu@HCl NPs with sizes of less than 100 nm, in addition to the formation of nanoparticles with diameters less than 10 nm **(Figure 2A)**. In contrast, Cu@HCl-NH_2_ NPs exhibited larger sizes with widths of approximately 0.25 μm to 1 μm. The rod-like structures in the TEM images provide a view of nanoplatelets with a vertical orientation, confirming that their thickness is in the nanometer range. Additionally, the thin size of the nanoplatelets is also confirmed by the slight reduction in transmission of electrons through areas with overlapping nanoplatelets. The difference in size between Cu@HCl NPs and Cu@HCl-NH_2_ NPs can be attributed to an effect of diamine on crystallization as the nanoplatelets are formed. The formation of the nanoplatelets on the copper TEM grids was further confirmed by scanning electron microscopy conducted on the TEM grids **(Figure 2B, Figure S1)**. Some nanoplatelets have either moved from the copper portions of the TEM grid to the carbon surface, or have newly formed on the carbon surface from copper ions that have diffused into the solution from the copper surfaces. To confirm the composition of the nanoplatelets, energy dispersive spectroscopy (EDS) was conducted on Cu@HCl-NH_2_ NPs at various locations on the carbon portion of the TEM grid **(Figure S2)**. EDS confirmed the presence of copper in these nanoplatelets, and also identified the presence of chlorine atoms, suggesting the nanoplatelets are likely composed of copper chloride. The presence of carbon and oxygen can be attributed to the carbon and oxygen atoms in the carbon grid coating, as well as the atoms in the diamine molecules.

**Figure 2.**
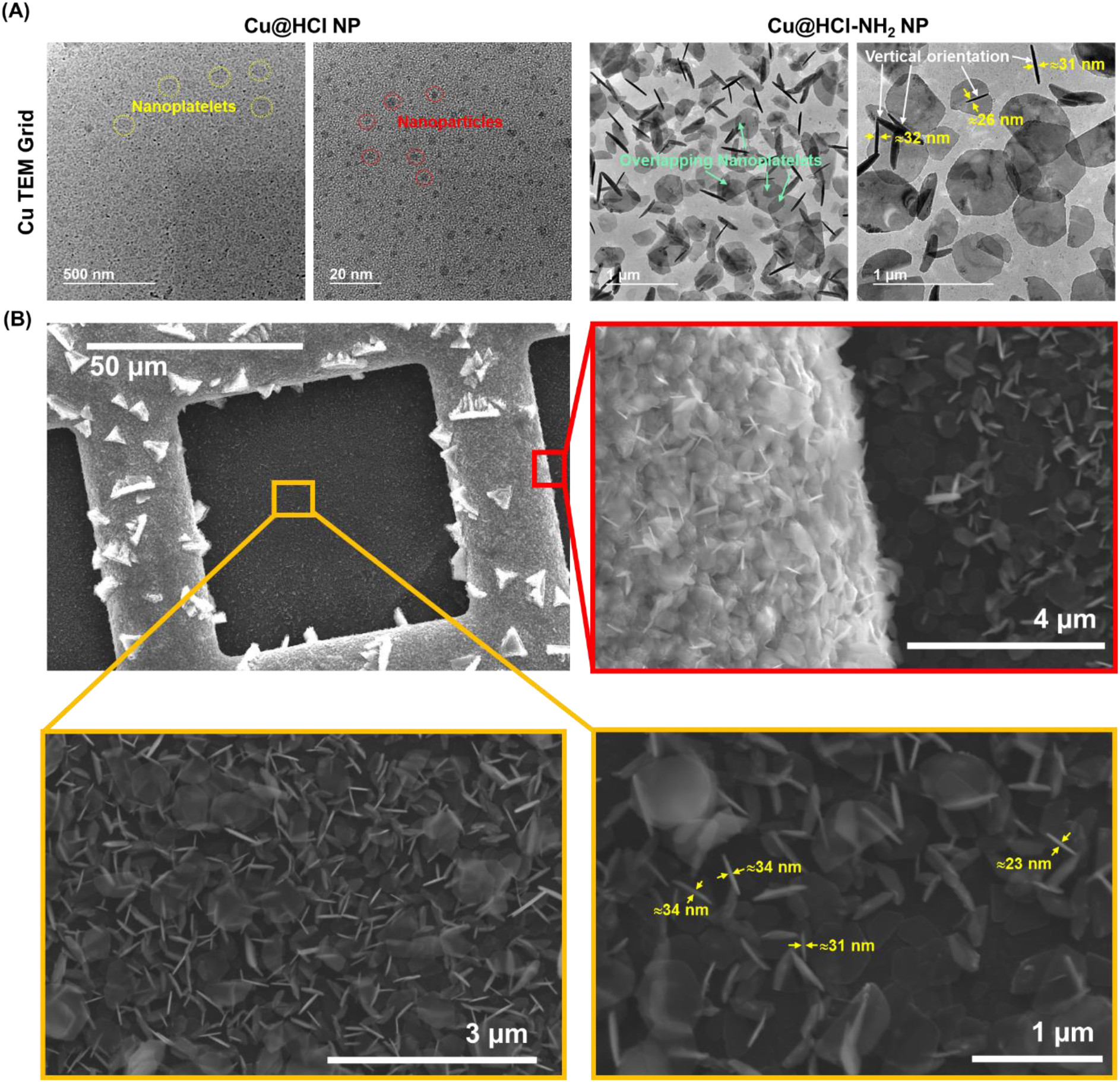
Copper chloride nanoplatelets formed on copper TEM grids. **(A)** TEM images of Cu@HCl NPs (left) and Cu@HCl-NH_2_ NPs (right), formed directly on the copper side of TEM grids (The contrast of the nanoplatelets is less than the contrast from the nanoparticles because the nanoplatelets are very thin). Nanostructures formed using the HCl solution alone include small nanoplatelets and nanoparticles, while the HCl and diamine solution results in the formation of hexagonal nanoplatelets. Rod-like structures are nanoplatelets that are oriented vertically rather than horizontally **(B)** SEM image of copper grid containing Cu@HCl-NH_2_ NPs. Nanoplatelets are found both on the copper portion of the grid and on the carbon portion of the grid, indicating the nanoplatelets can either move from the copper surface onto the carbon coating during the synthesis process, or form there from copper ions that have diffused from the copper surfaces.

Nanoplatelet precursor solutions were then deposited on copper tape surfaces to form nanoplatelets directly on the copper tape. Scanning electron microscopy images of the tape surfaces reveal the formation of Cu@HCl NPs with widths on the order of micrometers and thicknesses on the order of nanometers, as well as Cu@HCl-NH_2_ NPs that are wider than the Cu@HCl NPs **(Figure 3)**. The increase in Cu@HCl-NH_2_ NP widths can again be attributed to a role of diamine in the nanoplatelet crystallization process. XRD spectra reveal the presence of new peaks in 2θ ranges of 10° to 40° and 55° to 60° for Cu@HCl NPs and Cu@HCl-NH_2_ NPs, in addition to the peaks in the 2θ range of 40° to 55° that are from the Cu tape itself. The new peaks that resulted from nanoplatelet formation had some alignment with peaks in the XRD spectra of copper chloride, further suggesting that the nanoplatelets are composed of copper chloride crystals. However, SEM images of commercially obtained Cu(II) chloride powder revealed much larger and irregularly-shaped crystals **(Figure S3)**, indicating that the nanoplatelets that have been formed on the copper surfaces do not reflect the typical morphology of copper chloride powders, and are instead more ordered in structure. It is important to note that the XRD peaks resulting from copper tape cannot be separated from the signal resulting from the nanoplatelets, as the nanoplatelets are formed directly on copper tape. FT-IR spectra of Cu@HCl NPs and Cu@HCl-NH_2_ NPs reveal many spectral similarities to that of Cu tape, with the addition of multiple sharp peaks in 800 cm^−1^ to 1250 cm^−1^ range. Peaks at approximately 1580 cm^−1^, 1263 cm^−1^, and 1116 cm^−1^ are observed for Cu tape, Cu@HCl NPs, and Cu@HCl-NH_2_ NPs. Peaks at approximately 988 cm^−1^, 921 cm^−1^, and 834 cm^−1^ are observed for Cu@HCl NPs and Cu@HCl-NH_2_ NPs. A peak at approximately 1060 cm^−1^ is observed for only the Cu@HCl-NH_2_ NPs. These results suggest high similarity in composition between the Cu@HCl NPs, and Cu@HCl-NH_2_ NPs. As with the XRD spectra, it is important to note that the FTIR signal resulting from copper tape cannot be separated from the signal resulting from the nanoplatelets, as the nanoplatelets are formed directly on copper tape. Despite these limitations, these results suggest that the nanoplatelets are composed of copper chloride, although it cannot be said with certainty whether they are Cu(I) chloride or Cu(II) chloride. The ability for nanoplatelets to be synthesized on other copper surfaces such as copper electrodes and copper granules was also confirmed with SEM **(Figures S4, S5)**.

**Figure 3.**
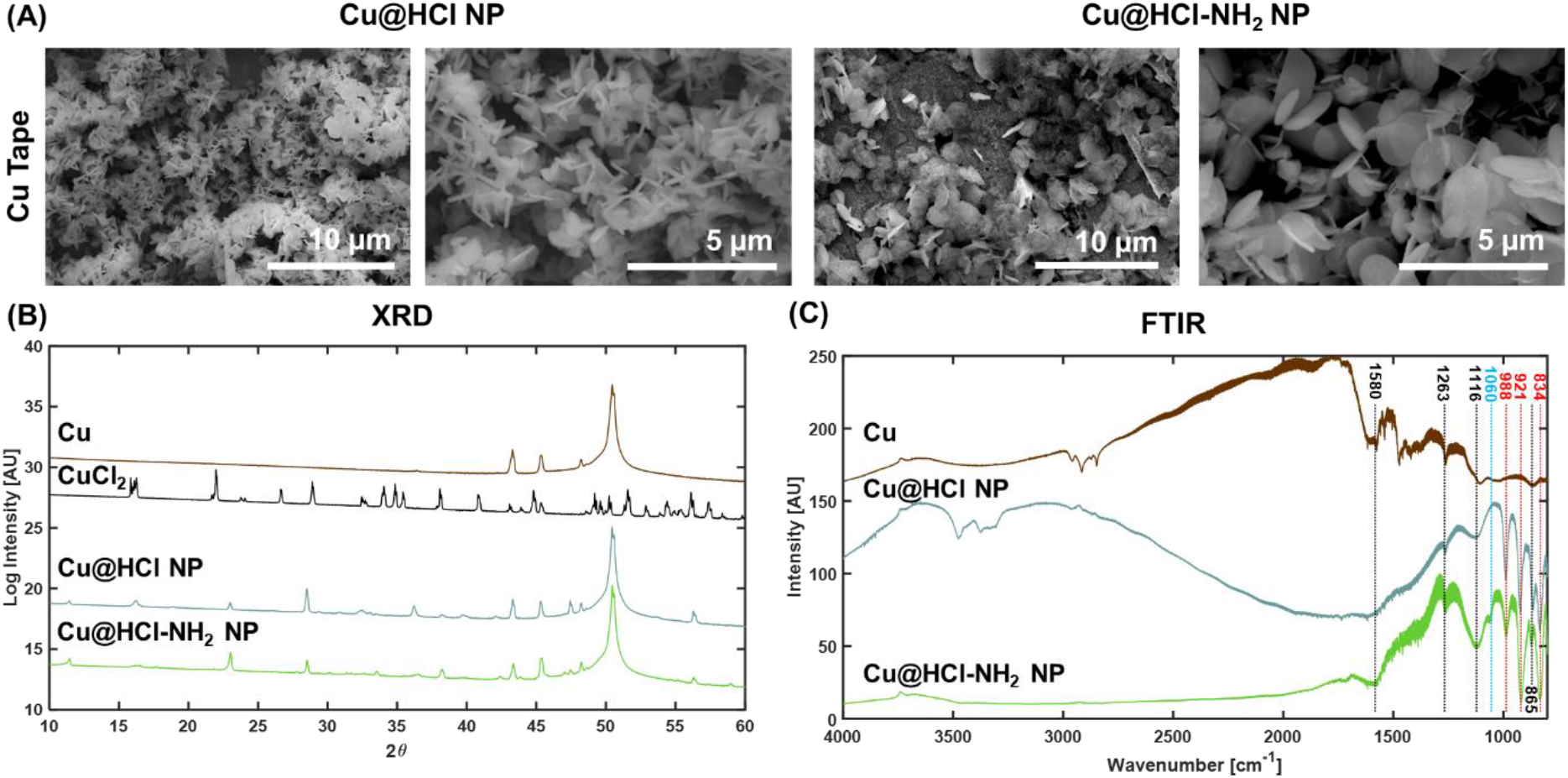
Characterization of copper chloride nanoplatelets formed on copper tape. **(A)** SEM images of nanoplatelets formed directly on copper tape: Cu@HCl NPs (left) and Cu@HCl-NH_2_ NPs (right). **(B)** XRD spectra of copper tape, copper (II) chloride powder, nanoplatelets formed directly on copper tape using the HCl solution, and nanoplatelets formed directly on copper tape using the HCl and diamine solution. **(C)** FTIR spectra of copper tape, Cu@HCl NPs, and Cu@HCl-NH_2_ NPs formed directly on copper tape. Black lines mark peaks present in Cu, Cu@HCl NPs, and Cu@HCl-NH_2_ NPs; Red lines mark peaks present in both Cu@HCl NPs and Cu@HCl-NH_2_ NPs but not in Cu; Blue line marks peak present in only Cu@HCl-NH_2_ NPs.

### Nanoplatelets inhibit *E. coli* and MDR *E. coli* bacterial growth

To assess the antibacterial properties of the surfaces with copper nanoplatelets compared to those without, an experimental setup was created to enable continuous contact between bacterial suspensions with those surfaces. Cu tape, Cu tape on which Cu@HCl NPs were synthesized, and Cu tape on which Cu@HCl-NH_2_ NPs were synthesized were adhered to the sides of 24-well plates **(Figure 4B)**. Bacterial suspensions were then added to the wells, and optical density at 600 nm (OD600) values representing bacterial concentration were measured over time. The initial color of the surfaces with Cu@HCl-NH_2_ NPs was observed to be bright green, while those with Cu@HCl NPs exhibited a lighter green color **(Figure 4A)**. With increasing exposure time to the bacterial solutions, a change in color of both the nanoplatelet tape samples and the solution was observed **(Figure 4A-B)**.

**Figure 4.**
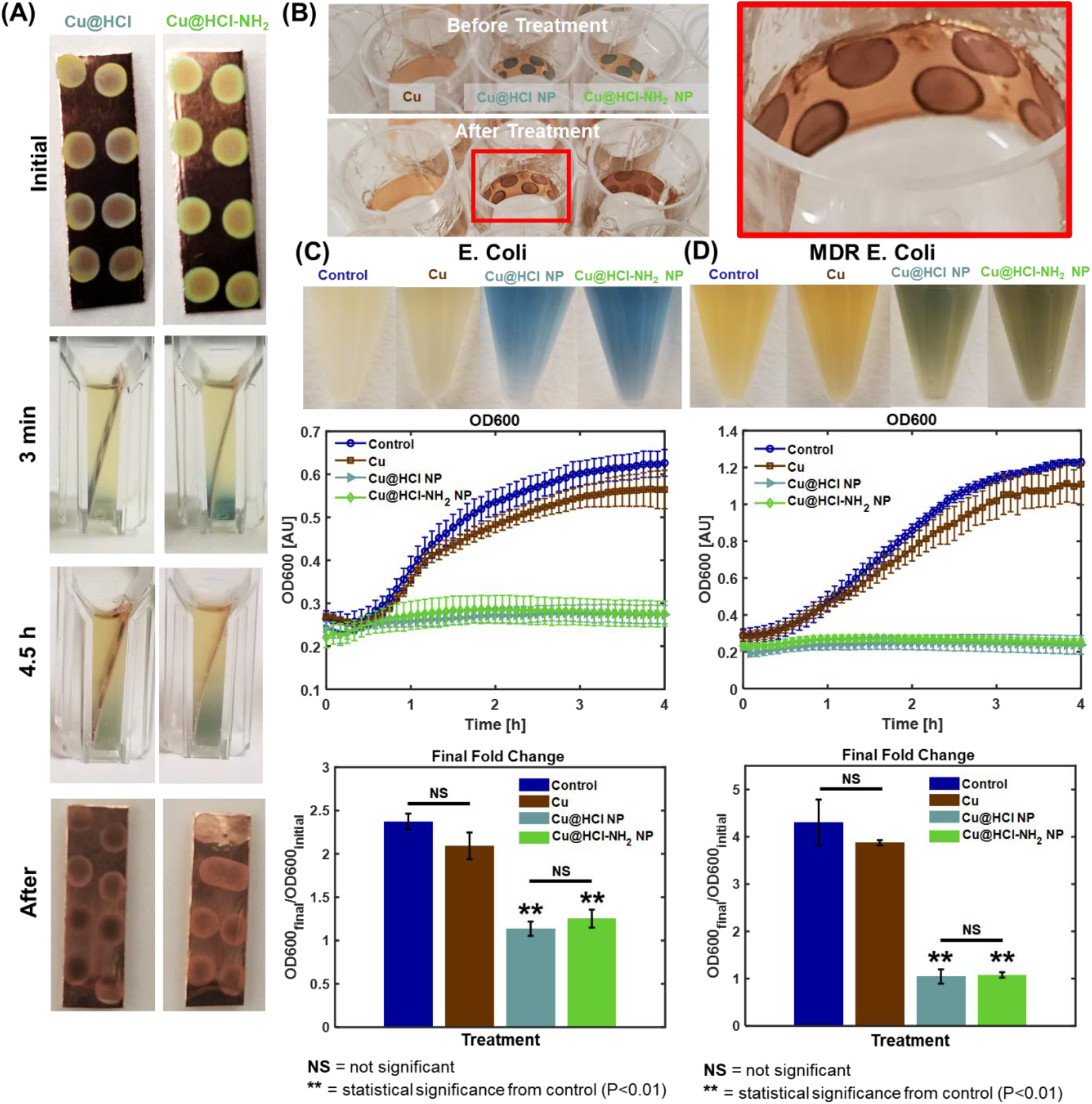
Inhibition of bacterial growth by nanoplatelets. **(A)** Nanoplatelet copper tape samples before and after exposure to bacteria. A visible color change occurs as a result of nanoplatelet dissolution into the bacterial broth. **(B)** Samples adhered to wells of 24-well plates. Top panel shows samples before exposure to bacteria, while bottom panel shows samples after exposure to bacteria. **(C)** Growth inhibition of *E. coli* in the presence of nanoplatelets. Nanoplatelets lead to almost complete inhibition of bacterial growth. Values represent averages and error bars represent standard deviations (n=8 for control, n=4 for treatment groups). **\*\*** Denotes statistical significance from control, with P<0.01. NS=not significant. Control samples are untreated bacterial solutions. **(D)** Growth inhibition of MDR *E. coli* in the presence of nanoplatelets. Nanoplatelets lead to almost complete inhibition of bacterial growth. Values represent averages and error bars represent standard deviations (n=8 for control, n=4 for treatment groups). **\*\*** Denotes statistical significance from control, with P<0.01. NS=not significant. Control samples are untreated bacterial solutions.

Changes in bacterial suspension color as a result of nanoplatelet treatment are apparent in both *E. coli* and MDR *E. coli* **(Figure 4C and 4D)**. The exposure of the broth to nanoplatelet-covered surfaces without bacteria results in the appearance of a UV-visible absorbance peak at 600 nm **(Figure S6)**. To account for the increase in absorbance at 600 nm due to copper tape with and without nanoplatelets in the absence of bacteria, the OD600 values for broth exposed to Cu tape and Cu tape with nanoplateletes were collected concurrently with, and subtracted from, the OD600 values for the bacteria-containing broth exposed to Cu tape and tape with nanoplatelet samples. In the presence of bacteria, the copper tape led to a smaller increase in OD600 values than when no tape was present at all, indicating a small amount of bacterial growth inhibition. However, the inhibition caused by the surfaces with Cu@HCl NPs and Cu@HCl-NH_2_ NPs represented a marked improvement over this. The OD600 values after 4 h of bacterial treatment were found to remain almost entirely unchanged for the Cu@HCl NPs and Cu@HCl-NH_2_ NPs, indicating almost complete inhibition of bacterial growth. This is also apparent in photographs of the wellplates after 4 h of treatment, with controls and bacterial suspensions exposed to copper tape having a greater opacity than suspensions exposed to copper tape with nanoplatelets **(Figure S7)**. To ensure that the observed bacterial growth inhibition was not caused by the HCl or HCl-NH_2_ solutions themselves, experiments were repeated with MDR *E. coli* exposure to liquid HCl solution or HCl-NH_2_ solution at equivalent concentrations to those used in forming the nanoplatelets **(Figure S8)**. The results reveal that the solutions themselves have little to no effect on OD600 values, confirming that the observed bacterial growth inhibition is a result of the presence of the nanoparticles on the copper surfaces. The mechanism of growth inhibition is not entirely clear. Considering the toxicity of copper ions,^34,35^ the inhibition likely results from the fact that the nanoplatelets increase the surface area from which copper ions can dissolve into solution and cause the growth inhibition.

### Scanning electron microscopy reveals bacterial damage as a result of nanoplatelet treatments

Scanning electron microscopy images were taken of control MDR *E. coli* samples, as well as those treated with Cu, Cu@HCl NPs, and Cu@HCl-NH_2_ NPs for 4 h **(Figure 5A)**. Control samples and Cu-treated samples retain a regular rod-like bacterial morphology, while samples exposed to nanoplatelet-covered copper surfaces have altered morphologies. In particular, exposure to those surfaces appears to lead to a wilting and flattening of bacterial samples. This is likely a result of an osmotic imbalance between the concentration of copper ions in the broth and in the bacterial cells. As copper ions are released from the dissolution of nanoplatelets into the bacterial suspension, a high concentration of copper ions would accumulate outside the bacterial cells and lead to hypertonic conditions. In response, the bacterial cells release water in an attempt to regain an equilibrium copper concentration between the bacteria and its environment, leading to the shriveled appearance in cells exposed to platelet-covered surfaces.

**Figure 5.**
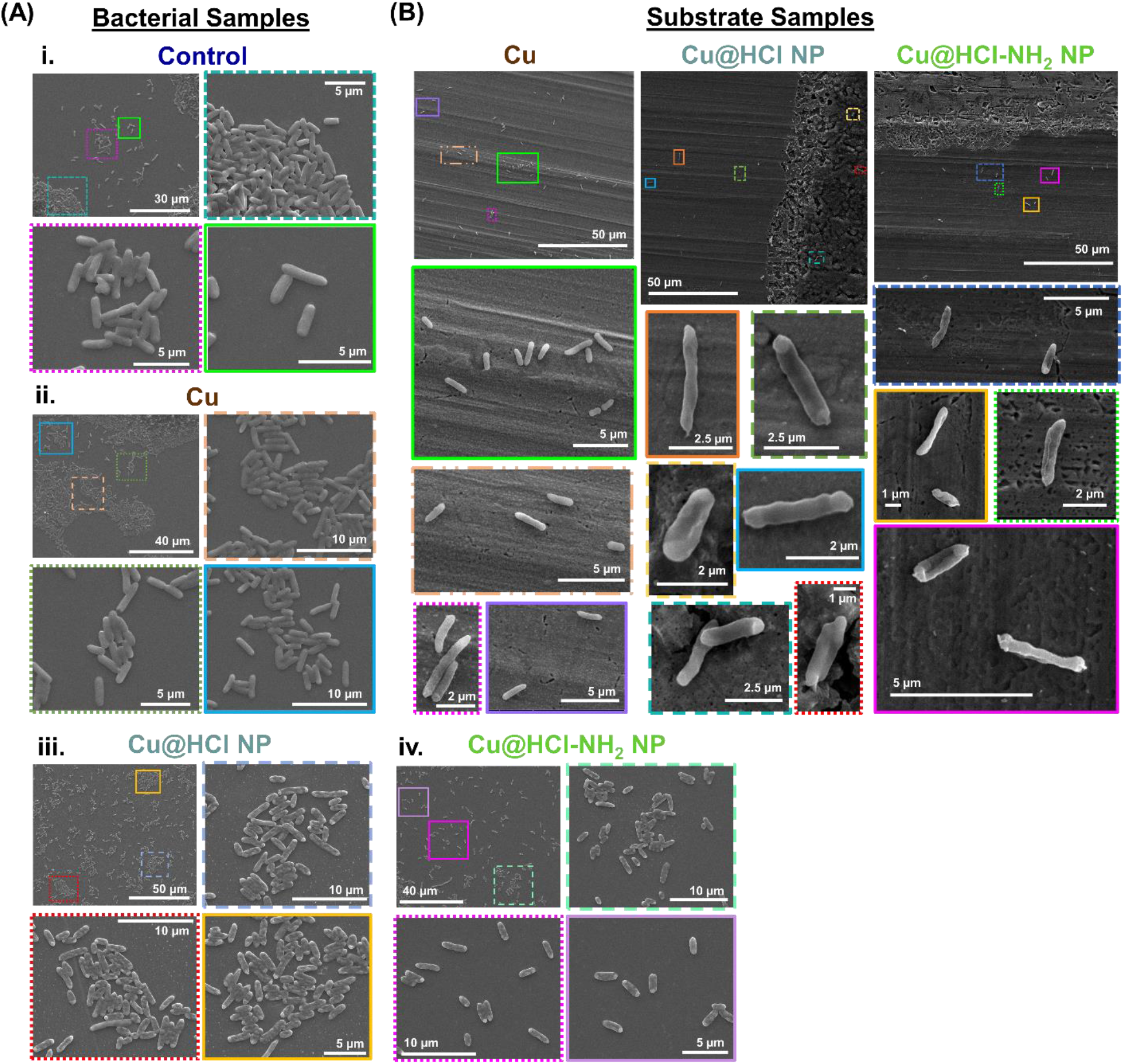
Scanning Electron Microscopy images of MDR *E. coli* treated with nanoplatelets. **(A)** Bacteria samples collected after 4 h of treatment. Insets depict higher magnification images of boxed regions. Untreated samples (controls) or samples treated with Cu tape alone exhibit regular bacterial morphologies **(i-ii)**. Samples treated with Cu@HCl NPs or Cu@HCl-NH_2_ NPs exhibit signs of bacterial damage, including a flattened and shriveled appearance **(iii-iv)**. **(B)** Bacteria on copper substrates after treatment. Insets depict higher magnification images of boxed regions. Bacteria treated with copper alone exhibit relatively normal morphologies, and exhibit some adhesion to the copper surface. Bacteria treated with nanoplatelets have minimal adhesion to the surface, and exhibit signs of damage.

Scanning electron microscopy images taken of the tape substrates with which MDR *E. coli* samples were treated (Cu, Cu@HCl NPs, and Cu@HCl-NH_2_ NPs) demonstrate a low adhesion between the bacterial cells and the copper substrates **(Figure 5B)**. Furthermore, bacteria on the Cu substrate samples appear to retain their normal rod-like morphology, while those on the nanoplatelet surfaces have a shriveled appearance, further confirming the negative effects of the presence of nanoplatelets on bacterial cell morphology.

### Exposure to nanoplatelet-covered copper surfaces reduced bacterial viability and induced DNA damage

Multiple studies were conducted to determine the mechanism by which the presence of nanoplatelets inhibit bacterial growth. Although OD600 measurements can provide information on the concentration of bacteria, they do not provide a direct measure of cell viability. To assess the viability of treated cells, live-dead staining of MDR *E. coli* was conducted using a commercial green stain for live cell indicator (live cell indicator) and propidium iodide (dead cell indicator), where a lower ratio of green to red fluorescence would indicate lower viability **(Figure 6A)**. These studies demonstrated that the presence of nanoplatelet-covered copper surfaces not only inhibit bacterial cell division, but also reduced bacterial cell viability within only 4 h of exposure. This result is consistent with copper ion toxicity observed by others.^34^

**Figure 6.**
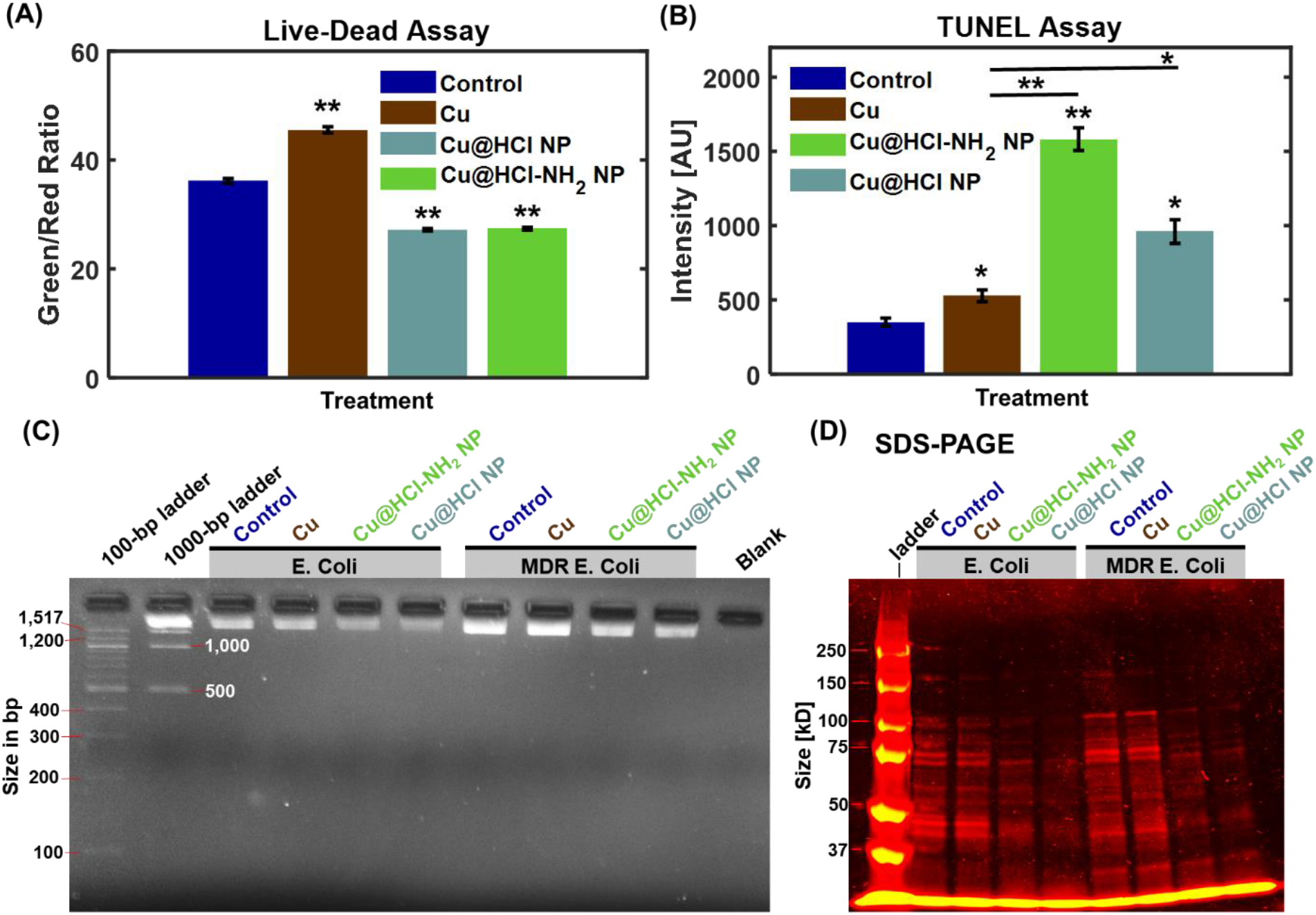
Exploration of mechanisms behind bacterial growth inhibition by nanoplatelets. **(A)** Live-Dead assay conducted on MDR *E. coli* treated with samples for 4 h. A higher Green/Red ratio indicates greater cell viability. A reduction in cell viability is caused by the nanoplatelet treatments. Values represent averages and error bars represent standard deviations across three technical replicates. **\*\*** indicates statistical significance with P<0.01. **(B)** TUNEL assay conducted on MDR *E. coli* treated with samples for 4 h. A higher intensity indicates a greater level of DNA damage. Nanoplatelets induce greater DNA damage than Cu alone. Values represent averages and error bars represent standard deviations across three technical replicates. Asterisks over individual bars indicate statistical significance with respect to the control. **\*\*** indicates statistical significance with P<0.01, **\*** indicates statistical significance with P<0.05. **(C)** Gel electrophoresis results for genomic DNA extracted from *E. coli* and MDR *E. coli* treated with samples for 4 h. Reductions in band intensities for Cu@HCl NP-treated and Cu@HCl-NH_2_ NP-treated samples confirm DNA damage induced by the nanoplatelets. These differences are apparent in both *E. coli* and MDR *E. coli*. **(D)** SDS-PAGE results for proteins extracted from *E. coli* and MDR *E. coli* treated with samples for 4 h. Nanoplatelets lead to reductions in band intensities, and in some cases complete removal of some protein bands, indicating that they have changes in protein expression. These changes are apparent in both *E. coli* than in MDR *E. coli*.

Since cell division appeared to be affected by the presence of nanoplatelet-covered copper surfaces, terminal deoxynucleotidyl transferase dUTP nick end labeling (TUNEL) assays were conducted to identify DNA damage in the bacterial samples **(Figure 6B)**. In TUNEL assays, fragmented DNA is labelled with a fluorescent dye, and the presence of greater DNA fragmentation leads to a greater fluorescent intensity. Cu alone was found to lead to some DNA fragmentation, as can be expected from the antibacterial properties of Cu. However, the presence of Cu@HCl NPs and Cu@HCl-NH_2_ NPs led to greater DNA fragmentation than the Cu treatment alone, confirming that the nanoplatelets provide a mechanism leading to a more potent antibacterial function. To further confirm this, gel electrophoresis was conducted on genomic DNA extracted from bacterial control samples, as well as bacterial samples treated with Cu, Cu@HCl NPs, and Cu@HCl-NH_2_ NPs **(Figure 6C)**. Exposure to nanoplatelet-covered surfaces reduced DNA band intensity for both *E. coli* and MDR *E. coli*, while untreated control samples and samples treated with Cu surfaces alone exhibited high band intensities, further confirming the role of nanoplatelets in preventing bacterial cell division by inducing DNA damage. To determine whether the nanoplatelets generated reactive oxygen species (ROS) that led to the DNA damage, ROS generation assays were conducted using CM-H_2_DCFDA, a fluorescent ROS indicator that has high retention in cells, enabling long-term studies. CM-H_2_DCFDA was loaded into MDR *E. coli* samples, and fluorescence intensity of the bacterial samples was monitored every 5 min for 4 h, with a greater fluorescence intensity corresponding to a higher presence of ROS. Although Cu alone was found to lead to some ROS generation near the beginning of the treatment, the presence of nanoplatelets on the copper surface was found to lead to lower fluorescence intensity than the control **(Figure S9)**. This indicates that the nanoplatelets may in fact be acting as ROS scavengers rather than inducing ROS generation. To ensure that this observation was not a result of CM-H_2_DCFDA photobleaching with time, the experiment was repeated with bacterial samples protected from light during the experiment, and measurements collected only after 4 h of treatment **(Figure S10)**. This further confirmed that ROS generation is not the mechanism by which nanoplatelet-induced DNA fragmentation occurred. Our results are consistent with those of Park et al., who found that copper ions do not increase hydroxyl radical production and may even reduce superoxide levels^35^.

### Exposure to nanoplatelet-covered copper surfaces led to changes in bacterial protein expression

In addition to evaluating DNA damage caused by the presence of nanoplatelets, further studies were conducted to evaluate the effects of nanoplatelet-covered copper surfaces on bacterial protein expression. Water-soluble proteins were extracted from untreated control samples, as well as bacteria exposed to copper surfaces, Cu@HCl NPs, and Cu@HCl-NH_2_ NPs. SDS PAGE was conducted on these protein samples **(Figure 6D)**, and demonstrates a clear difference in protein expression for nanoplatelet-treated bacteria compared to Cu-treated or untreated bacteria. These changes in protein expression are apparent for both *E. coli* and MDR *E. coli*. Furthermore, exposure to copper surfaces without nanoplatelets appeared to lead to little change in protein expression in either type of bacteria.

To examine this in greater detail, proteomic analysis was carried out to identify individual proteins present in the protein population extracted from the bacteria. Heatmaps were created using the −10logP values for all detected proteins for *E. coli* and MDR *E. coli*, where a higher −10logP value corresponds to a greater confidence in the identification of the proteins. In this work, we have divided −10logP values into 3 ranges: values greater than 75 are considered identification of the proteins with high confidence, values between 45 and 75 are considered identification of the proteins with acceptable confidence, and values below 45 are considered proteins that cannot be confidently identified in the samples. A list of all protein IDs and their corresponding protein Accession IDs and descriptions are provided in **Table S1 and Table S2**.

**Proteomic Analysis of MDR *E. coli***. There were fewer identified proteins in MDR *E. coli* samples exposed to copper surfaces without nanoplatelets (450 proteins), Cu@HCl-covered copper surfaces (130 proteins), and Cu@HCl-NH_2_-covered copper surfaces (182 proteins) than in control samples that were not exposed to either surfaces (577 proteins). This confirms SDS PAGE observations which illustrated that exposure to nanoplatelet-covered Cu surfaces had a major impact on protein expression.

A heatmap of identified proteins in the MDR *E. coli* samples is provided in **Figure 7A**. Three proteins related to cell division were identified in the control sample: 89 (P0A9A6|FTSZ_ECOLI, Cell division protein FtsZ), 630 (P0AF36|ZAPB_ECOLI, Cell division protein ZapB), and 734 (P45955|CPOB_ECOLI, Cell division coordinator CpoB). Protein 89 was identified with lower −10logP values in bacterial samples treated with Cu, Cu@HCl NPs, or Cu@HCl-NH_2_ NPs, and protein 734 could not be confidently identified in these samples. A down-regulation of cell division proteins as a result of treatment can be directly related to the lack of cell division observed during the OD600 measurements.

**Figure 7.**
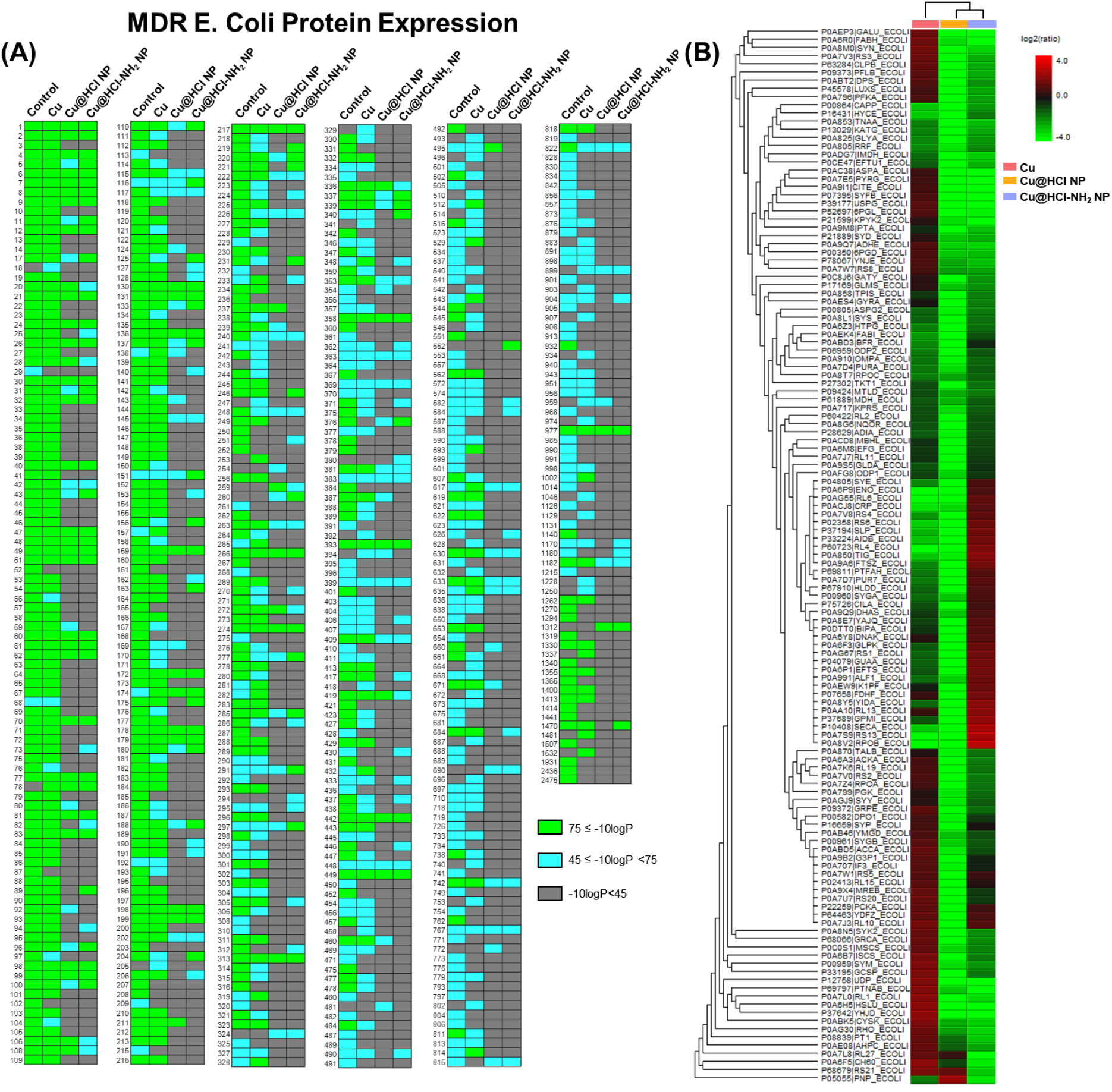
Proteomics analysis of MDR *E. coli*. **(A)** Heat maps classifying the −10logP values of each protein in each sample. Data represent a compilation of three technical replicates taken from each sample. **(B)** Hierarchical clustering of ratios between protein relative expression in each sample and protein relative expression in the control sample. Data represent the most data-rich technical replicate of three technical replicates for each sample.

Additionally, protein 78 (Q59385|COPA_ECOLI, Copper-exporting P-type ATPase) was identified in MDR *E. coli* treated with either Cu or nanoplatelets, but not in the control sample. Protein 458 (P36649|CUEO_ECOLI, blue copper oxidase), a protein believed to be involved in the detoxification of copper, was also identified in Cu-treated samples, but not in the control or the nanoplatelet-treated samples. Thus, the DNA damage caused by exposure to nanoplatelet-coated copper surfaces may have prevented the eventual synthesis of proteins that would provide the MDR *E. coli* with copper tolerance. In contrast, exposure to uncoated Cu surfaces may have led to the synthesis of proteins that would increase the copper tolerance of MDR *E. coli*.

We further examined proteins related to DNA. Protein 58 (P0AES4|GYRA_ECOLI, DNA gyrase subunit A), which is known to be involved in ATP-dependent breakage, passage and rejoining of double-stranded DNA, and therefore plays a role in its transcription, repair, and replication, is present in the control sample and Cu-treated sample, but not in the nanoplatelet-treated samples. Protein 90 (P00582|DPO1_ECOLI, DNA polymerase I) is also present in the control samples and samples exposed to uncoated copper surfaces, but not in those exposed to nanoplatelet-coated copper surfaces. This protein exhibits polymerase and exonuclease behaviors. Protein 114 (P0AES6|GYRB_ECOLI, DNA gyrase subunit B), which relaxes negatively supercoiled DNA, is identified with less confidence in the sample exposed to Cu surfaces than the control samples, and is not confidently identified in the samples exposed to nanoplatelet-covered Cu surfaces. Proteins 410 (P06612|TOP1_ECOLI, DNA topoisomerase 1) and 1215 (P12295|UNG_ECOLI, Uracil-DNA glycosylase) could not be confidently identified in any of the treated samples, despite being identified in the control sample. These proteins are responsible for ATP-independent breakage of single-stranded DNA and releasing uracil residues respectively. Finally, protein 529 (P0ABE2|BOLA_ECOLI, DNA-binding transcriptional regulator BolA), which is known to have an impact on cell morphology, cell growth, and cell division, is identified in the control and Cu-exposed samples, but not confidently identified in the samples exposed to nanoplatelet-coated surfaces. Altogether, these results confirm that the exposure to nanoplatelet-covered surfaces had an impact on MDR *E. coli* cell division, response to copper, and DNA replication **(Table 1)**.

**Table 1.**
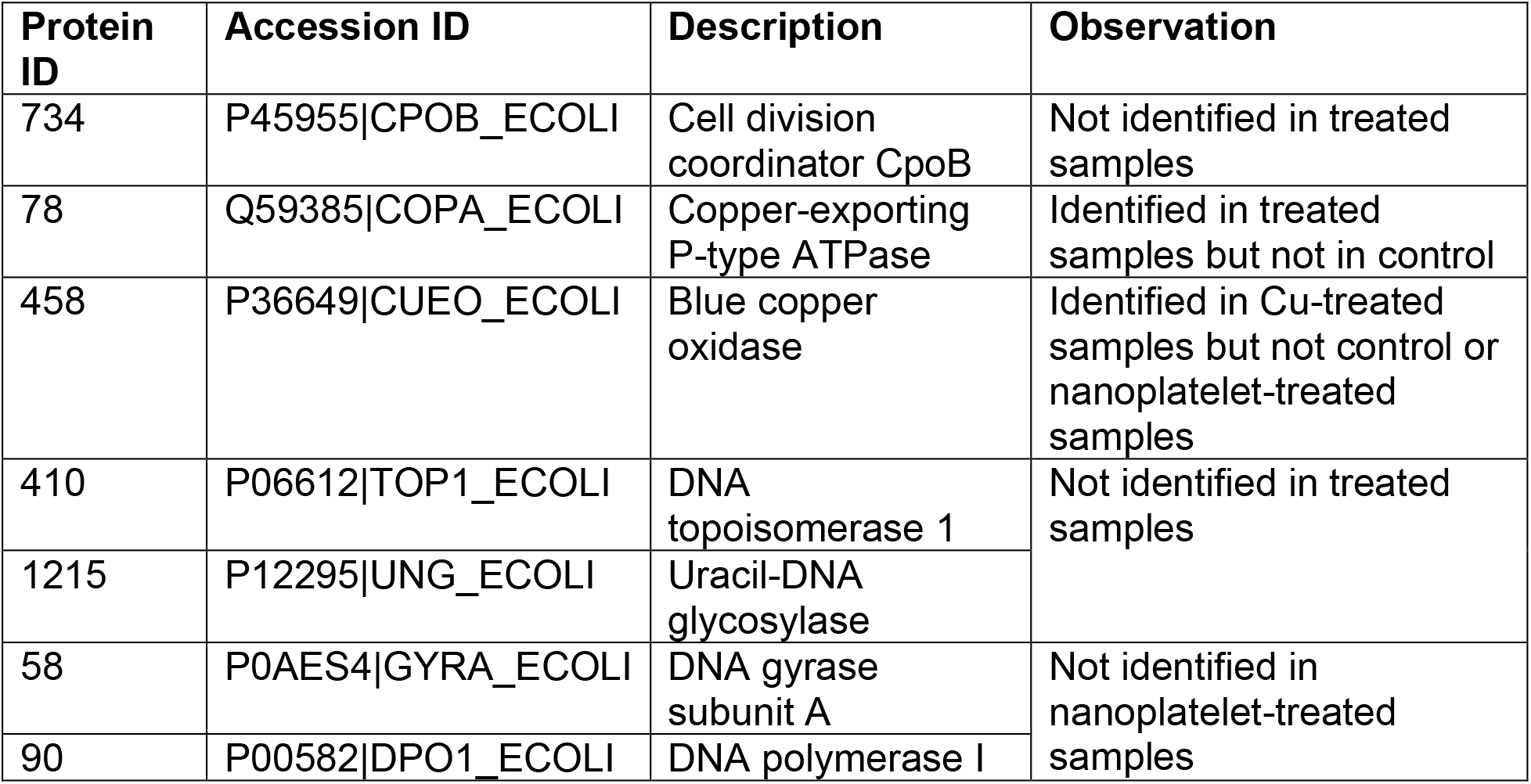
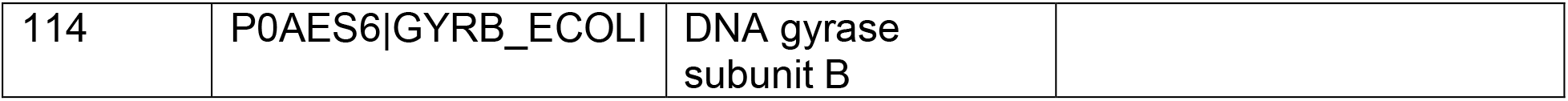
Proteins of interest in MDR *E. coli* samples.

To develop an understanding of each sample’s protein expression relative to the control sample, the ratio of each protein’s ion counts to the total ion counts for each sample were calculated. This relative expression ratio was then divided by the relative expression ratio of the same proteins in the control sample. This enabled a comparison between the relative abundance of different proteins in the treated samples compared to the untreated control. Hierarchical clustering of these protein ratios identified similarities between samples exposed to nanoplatelet-covered surfaces compared to the samples exposed to copper surfaces without nanoplatelets **(Figure 7B)**. In addition, the presence of nanoplatelets generally led to decreased expression of many proteins, while the absence of nanoplatelets on copper surfaces led to increased expression of some proteins critical to survival. For example, Cu-treated samples had an increased relative expression (compared to the control) of P0ABT2|DPS_ECOLI, which is responsible for protecting the bacterial cells from copper ion toxicity.

**Proteomic Analysis of *E. coli***. As for MDR *E. coli*, there are fewer proteins in the *E. coli* samples treated with Cu (377 proteins), Cu@HCl NPs (178 proteins), and Cu@HCl-NH_2_ NPs (110 proteins) compared to the untreated control (611 proteins). This further confirms the *E. coli* SDS PAGE observations which illustrated that the presence of nanoplatelet had a major impact on protein expression.

A heatmap of identified proteins in the *E. coli* samples is provided in **Figure S11 A.** A number of proteins related to cell division were identified in the control sample. Protein 89 (P0A9A6|FTSZ_ECOLI, Cell division protein FtsZ), which is responsible for forming a contractile ring structure (Z ring) at the future cell division site, was identified with high confidence in both the control and Cu-treated samples, but was identified with lower confidence in Cu@HCl NP-treated copper surfaces and could not be confidently identified in sample exposed to Cu@HCl-NH_2_ NP-treated copper surfaces. Protein 900 (P0A734|MINE_ECOLI, Cell division topological specificity factor), which is responsible for ensuring the occurrence of cell division at the proper site, was identified in the control sample but not in the treated samples.

A number of proteins related to DNA were also altered in expression as a result of the exposure to copper surfaces. Protein 260 (P0ACF0|DBHA_ECOLI, DNA-binding protein HU-alpha), protein 265 (P0A800|RPOZ_ECOLI, DNA-directed RNA polymerase subunit omega), protein 272 (P0ACG1|STPA_ECOLI, DNA-binding protein StpA), protein 402 (P23909|MUTS_ECOLI, DNA mismatch repair protein MutS), protein 528 (P0ACB0|DNAB_ECOLI, Replicative DNA helicase), protein 837 (P21189|DPO2_ECOLI, DNA polymerase II), protein 1250 (P0ABS5|DNAG_ECOLI, DNA primase), and protein 1294 (P65556|YFCD_ECOLI, DNA primase) were all identified in the control sample but could not be confidently identified in any of the samples exposed to any of the copper surfaces (both coated and uncoated with nanoplatelets). Proteins 80 (P0A7Z4|RPOA_ECOLI, DNA-directed RNA polymerase subunit alpha), 85 (P0A9X4|MREB_ECOLI, DNA-directed RNA polymerase subunit omega), 86 (P04805|SYE_ECOLI, DNA-directed RNA polymerase subunit alpha), 90 (P00582|DPO1_ECOLI, DNA polymerase I), 114 (P0AES6|GYRB_ECOLI, DNA gyrase subunit B), and 529 (P0ABE2|BOLA_ECOLI, DNA-binding transcriptional regulator BolA) were all identified in the control and Cu-treated samples but not in the samples exposed to nanoplatelet-covered surfaces **(Table 2)**.

**Table 2.**
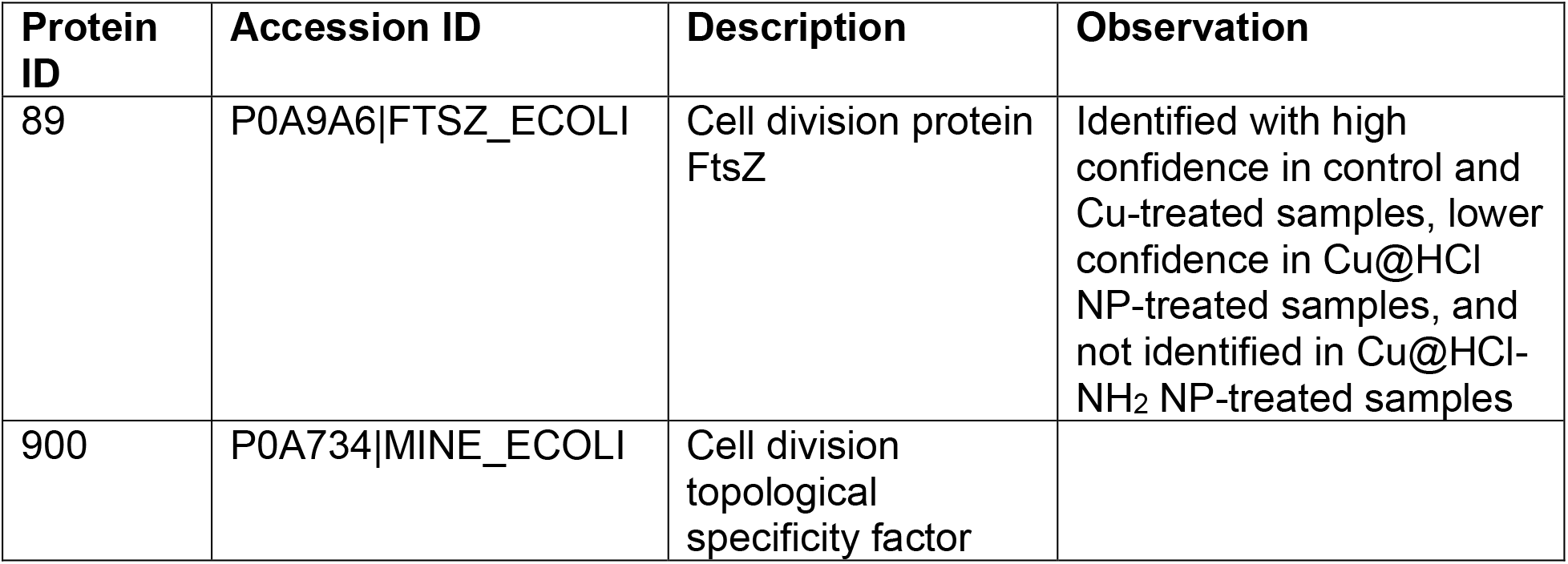
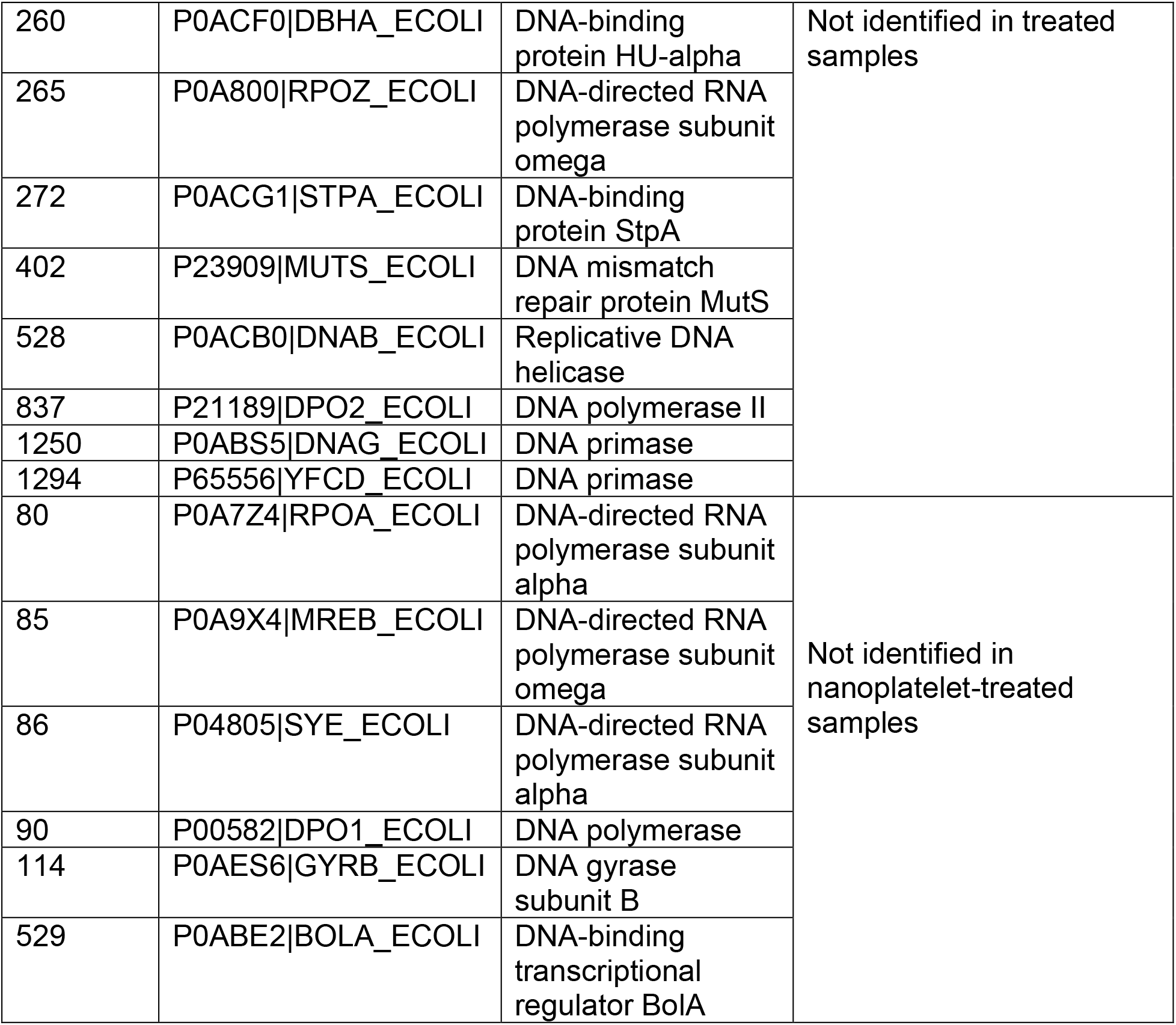
Proteins of interest in *E. coli* samples.

Hierarchical clustering of *E. coli* protein ratios identified similarities between the samples exposed to nanoplatelet-covered surfaces compared to the samples exposed to bare copper surfaces **(Figure S 11 B)**. As with MDR *E. coli*, exposure to nanoplatelets generally led to decreased expression of many proteins, while exposure to Cu surfaces without nanoplatelets led to increased expression of many proteins critical to survival. These findings further confirm the role of the presence of nanoplatelets in modulating protein expression.

### Nanostructures were also assembled directly on multiple other metal surfaces

To determine whether nanoplatelets could be assembled on other metal surfaces, the synthesis procedures were repeated by depositing the HCl solution or HCl and diamine solution onto pieces of tin, zinc, and cobalt and allowing the solution to dry. Scanning electron microscopy was then conducted on the samples **(Figure S12, Figure S13)**. These images revealed the successful formation of nanostructures on these various metal surfaces. Although the structures differed in shape from one metal type to another, these results suggest a simple aqueous solution containing acid can be applied to metal surfaces to obtain textured nanoscale patterns and structures on the surfaces.

## CONCLUSIONS

We have developed a simple method to assemble metal-based nanostructures directly on metal surfaces without the need for prior synthesis of nanoparticles or inclusion of any metals within the precursor solution. We demonstrated that nanoplatelets could be formed on a variety of copper surfaces and explored their potential for use in antibacterial surfaces. We found that the copper surfaces with nanoplatelets had improved antibacterial activity compared to copper surfaces without nanoplatelets, likely as a result of an increased surface area from which copper ions can dissolve. Copper surfaces covered with nanoplatelets also induced structural and morphological changes in bacterial cells, and led to changes in protein and DNA expression. We further demonstrated that expression of specific proteins related to cell division, copper toxicity, and DNA division were altered as a result of the presence of nanoplatelets. Last, we demonstrated that other nanostructures could be formed by depositing our simple aqueous diluted HCl solution onto various metal surfaces. Overall, this study provides a simple method by which metal-based surfaces, especially those containing copper, can be modified to incorporate 2D nanostructures.

## MATERIALS AND METHODS

### Materials

Copper granules, zinc strips, tin chips, and cobalt pieces were purchased from Chemistry Cabinet^‡^. Other metal samples were obtained in the form of a copper anode sheet and zinc anode sheet. 300 to 400 mesh carbon coated copper TEM grids were purchased from Ted Pella^‡^. Regular *E. coli* and multi-drug-resistant *E. coli* (ATCC BAA-201) were purchased from ATCC^‡^. Loctite Clear Silicone Waterproof Sealant^‡^ was used to adhere copper substrate samples to the wells of 24-well plates. DNA extraction was conducted using a ZymoBIOMICS DNA Miniprep Kit (Zymo Research)^‡^. Protein extraction was conducted using a Qproteome Bacterial Protein Prep Kit (Qiagen)^‡^, and protein digestions were conducted using an In-Solution Tryptic Digestion and Guanidination Kit (89895, ThermoFisher Scientific)^‡^. CM-H_2_DCFDA and the LIVE/DEAD BacLight Bacterial Viability Kit were purchased as kits from ThermoFisher Scientific^‡^. The Cell Meter TUNEL apoptosis assay kit was purchased from AAT Bioquest^‡^.

### Preparation of Nanoplatelet Precursor solution

*Solution for Cu@HCl NP:* 50 μL of 2N HCl was diluted with 1 mL of filtered water. *Solution for Cu@HCl-NH_2_ NP:* 3.5 μL 2 2’-(ethylenedioxy) bis(ethylamine) was diluted in 10 mL of filtered water. 50 μL of 2N HCl was added to 1 mL of this solution.

### Synthesis of Nanoplatelets

*On TEM Grids:* 2.5 μL of precursor solution was drop-cast onto the uncoated side of carbon-coated copper TEM grids. Samples were allowed to dry for 2 min, after which excess liquid was removed using filter paper. The TEM grid was then allowed to dry until the imaging was conducted.

*On Copper Tape and other metal substrates:* 3.5 μL of precursor solution was drop-cast onto the substrate and allowed to completely dry under atmospheric conditions. For bacterial experiments, eight 3.5 μL drops of precursor solution were deposited on each 2-cm copper sample.

### Scanning Electron Microscopy of Nanoplatelet Samples

Samples were imaged using a commercial environmental SEM at nominal acceleration voltages of 10 kV to 20 kV. Samples were not coated prior to imaging.

### FT-IR and XRD

FT-IR samples were prepared by adhering copper tape samples directly onto corner frosted FT-IR slides. FT-IR measurements were taken with a commercial FT-IR spectrometer. XRD was conducted on powder samples using a commercial XRD system. XRD spectra were collected for nanoplatelet samples directly on the Cu tape substrate.

### Bacterial Culture

*E. coli* was cultured in Luria-Bertani (LB) broth at 37 °C under aerobic conditions. MDR *E. coli* was cultured at 37 °C under aerobic conditions in Tryptic Soy Medium supplemented with 10 μg/ml Ceftazidime.

### Bacterial optical density experiments

Copper substrates or nanoplatelet samples were adhered to the wells of a 24-well plate using waterproof silicone glue. Two 2 cm sample strips were used for each well. The glue was allowed to completely dry before experiments were conducted. 2 mL of bacteria with an optical density of approximately 0.25 were added to each well of the 24-well plates. OD600 values were measured kinetically at 37 °C for 4 h in a plate reader.

### Bacterial optical density experiments using precursor solutions

2 mL of MDR *E. coli* bacteria with an optical density of approximately 0.25 were added to 12 wells of a 24-well plate. 2 mL of bacterial broth were added to the remaining 12 wells of a 24-well plate. For treatment groups, 56 μL of nanoplatelet precursor solution was added to each well. OD600 values were measured kinetically at 37 °C for 4 h in a plate reader.

### Scanning Electron Microscopy of substrates used to treat bacteria

2 mL of MDR *E. coli* with an initial optical density of approximately 0.25 was exposed to two 2 cm long copper tape samples or nanoplatelet samples for 4 h at 37 °C in an incubator-shaker. Following this, samples were fixed in a 2 % paraformaldehyde, 2.5 % glutaraldehyde solution for 10 min, and sequentially dehydrated in solutions of 25 %, 50 %, 75 %, 95 %, and 100 % ethanol for 10 min each. The samples were then allowed to dry under atmospheric conditions. Prior to imaging, samples were coated with platinum-palladium. Samples were imaged on a commercial SEM with a 10 kV nominal acceleration voltage.

### Scanning Electron Microscopy of bacteria after exposure to substrates

2 mL of MDR *E. coli* with an initial optical density of approximately 0.25 was exposed to two 2 cm long copper tape samples or nanoplatelet samples for 4 h at 37 °C in an incubator-shaker. Following this, the bacterial suspensions were centrifuged at 626 rad/s (5976 rpm), 39200 m/s^2^ (4000, × g) for 15 min, and resuspended in 0.5 mL 2% volume fraction paraformaldehyde/water solution, 2.5% volume fraction glutaraldehyde/water solution for 10 min. The suspensions were then centrifuged for 10 min and resuspended in 25% volume fraction ethanol/water solution for 10 min. Suspensions were then sequentially centrifuged (626 rad/s (5976 rpm), 39200 m/s^2^ (4000 × g), 5 min) and resuspended in 0.5 mL of 50%, 75%, and 95% volume fraction ethanol/water solutions, with samples retained in each solution for 5 min prior to centrifugation. Finally, the samples were in 250 μL 100% ethanol, and 50 μL of each sample was deposited onto a glass coverslip and allowed to dry under atmospheric conditions. Prior to imaging, samples were coated with platinum-palladium. Samples were imaged on a commercial SEM with a 10 kV nominal acceleration voltage.

### ROS Generation Assay

50 μg CM-H_2_DCFDA dye was dissolved in 8.65 μL dimethyl sulfoxide [DMSO], and 8 μL of this solution was added to 8 mL bacterial broth. 20 mL of bacteria was suspended at an OD600 value of 0.25. The bacteria was centrifuged at 700 rad/s (6682 rpm), 49000 m/s^2^ (5000 × g) for 5 min and then resuspended in the bacterial medium containing the CM-H_2_DCFDA dye. The bacteria was protected from light and incubated in a shaker-incubator at 37 °C for 5 min. The suspension was then centrifuged at 626 rad/s (5976 rpm), 39200 m/s^2^ (4000 × g) for 5 min, and the bacteria was resuspended in 8 ml fresh medium. The centrifugation process was repeated one more time, and the bacterial pellet was finally resuspended in 20 mL fresh medium. 1 mL of bacterial suspension was placed in each centrifuge tube, and the suspensions were treated with the copper tape or nanoplatelets, after which fluorescence measurements were collected in accordance with kit directions.

### Live-Dead Assay

2 mL of MDR *E. coli* with an initial optical density of approximately 0.25 was exposed to two 2 cm long copper tape samples or nanoplatelet samples for 4 h at 37 °C in an incubator-shaker. Following this, 0.75 mL of each sample was collected and centrifuged to form a bacterial cell pellet. The pellet was then resuspended in NaCl solution with a volume fraction of 0.85 %, and three 100 μL aliquots from each solution were added into the wells of a 96-well plate. 100 μL of live-dead solution (prepared according to manufacturer guidelines) was added to each well. Samples were incubated in the dark at room temperature for 15 min, after which fluorescence spectra were collected

### TUNEL Assay

2 mL of MDR *E. coli* with an initial optical density of approximately 0.25 was exposed to two 2 cm long copper tape samples or nanoplatelet samples for 4 h at 37 °C in an incubator-shaker. Following this, 0.75 mL of each sample was collected and centrifuged to form a bacterial cell pellet. Each pellet was then resuspended in 150 μL TUNEL solution (12.5 μL tunnelyte diluted in 1.25 mL reaction buffer) and placed on an incubator-shaker at 37 °C for 1 h. Samples were then pelleted and resuspended in 300 μL reaction buffer and three 100 μL aliquots (3 technical replicates) of each sample were pipetted into the wells of a 96-well plate. Fluorescence spectra were collected for each well (excitation 550 nm, emission range: 590 nm to 650 nm, gain of 150), and the peak emission intensity was averaged across all technical replicates for each sample.

### DNA Extraction

2 mL of MDR *E. coli* with an initial optical density of approximately 0.25 was exposed to two 2 cm long copper tape samples or nanoplatelet samples for 4 h at 37 °C in an incubator-shaker. DNA was extracted following the guidelines provided by the kit manufacturer. Samples were stored frozen at −20 °C until further use.

### DNA gel electrophoresis

Agarose gels were prepared by adding 2 g agarose to 100 mL Tris-Borate-EDTA (TBE) buffer and heating the solution in a microwave. Ethidium bromide was added into the solution, and the gel was poured into the mold and allowed to cool. 50 ng of each DNA sample was added to their respective wells, and the gel was run at 120 V for 75 min.

### Protein Extraction

2 mL of MDR *E. coli* with an initial optical density of approximately 0.25 was exposed to two 2 cm long copper tape samples or nanoplatelet samples for 4 h at 37 °C in an incubator-shaker. Water-soluble proteins were then extracted from bacteria following the guidelines in the obtained kit. Samples were stored frozen at −80 °C until further use.

### SDS PAGE

Protein concentration was determined and samples were diluted to a concentration of 1 μg/μL and run on a 7.5% pre-cast gel at 115 V for 48 min, after which they were stained and imaged on a gel imager.

### Statistical Analysis

Statistical analysis was conducted with two-tailed Student’s T-tests. Bonferroni corrections were used to determine adjusted P values in cases where multiple comparisons were conducted.

### Proteomics Analysis

Protein concentration was determined using a Nanodrop^‡^. Samples were diluted to a concentration of 1 μg/μL and a 10 μL aliquot was digested following the kit instructions. Samples were stored at −20 °C and submitted to the UMBC Molecular Characterization and Analysis Complex for label-free proteomic analysis.

## ACKNOWLEDGEMENTS

We gratefully acknowledge Dr. Priyanka Ray and Dr. Parikshit Moitra for their helpful comments and assistance with electrophoresis experiments, as well as Dr. Aaron Schwartz-Duval for his helpful comments. This project was partially funded through grants from the National Institutes of Health, Department of Defense, and University of Illinois. P. Fathi was supported by the National Physical Science Consortium and the National Institute of Standards & Technology through a National Physical Science Consortium (NPSC) graduate fellowship and by the Nadine Barrie Smith Memorial Fellowship from the Beckman Institute. Research reported in this publication was supported by the National Institute of Biomedical Imaging and Bioengineering of the National Institutes of Health under Award Number T32EB019944. This work was carried out in part in the Frederick Seitz Materials Research Laboratory Central Research Facilities, University of Illinois.

## Footnotes

‡ Commercial entities, equipment or materials may be identified in this document to describe an experimental procedure or concept adequately. Such identification is not intended to imply recommendation or endorsement by the National Institute of Standards and Technology, nor is it intended to imply that the entities, materials or equipment are necessarily the best available for the purpose.

