## Supporting Information for "*In Situ* Surface-Directed Assembly of 2D Metal Nanoplatelets for Drug-Free Treatment of Antibiotic-Resistant Bacteria"

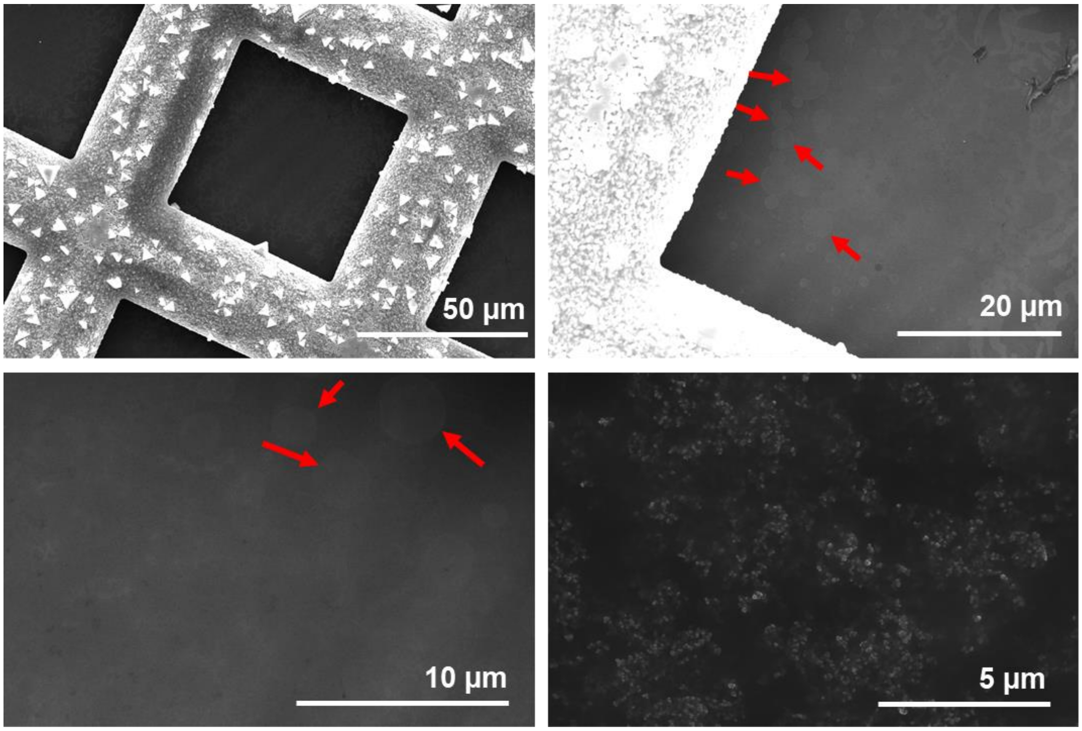


**Figure S1.** SEM image of copper grid containing Cu@HCl NPs. Nanoplatelets are found both on the copper portion of the grid and on the carbon portion of the grid, indicating the nanoplatelets can move from the copper surface onto the carbon coating during the synthesis process. The presence of small Cu@HCl nanoparticles was also observed.

**Location A**

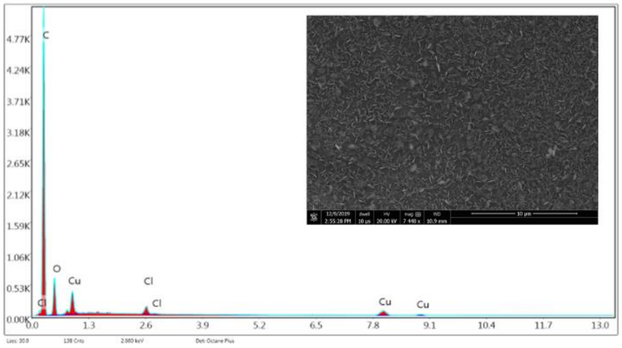


**Location B**

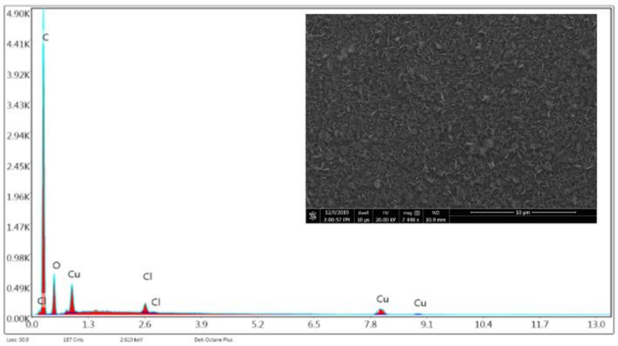


**Location C**

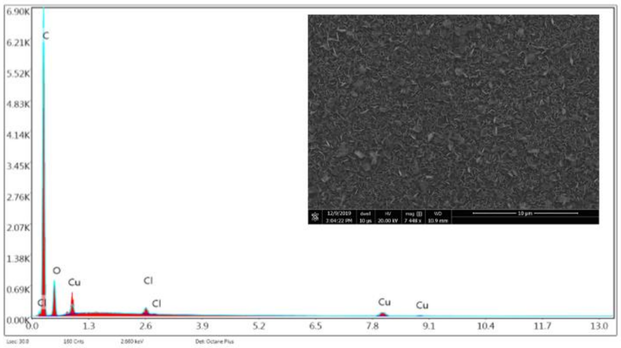


**Location A**

| Element | Weight % | Atomic % | Error % | Net Int. | K Ratio | Z | A | F |
|---|---|---|---|---|---|---|---|---|
| C K | 75.63 | 82.86 | 5.46 | 893.07 | 0.4724 | 1.0197 | 0.6125 | 1 |
| O K | 19.41 | 15.97 | 13.09 | 111.92 | 0.0252 | 0.9746 | 0.1332 | 1 |
| Cl K | 0.85 | 0.31 | 10.19 | 42.21 | 0.0071 | 0.8232 | 1.0038 | 1.0171 |
| Cu K | 4.11 | 0.85 | 8.4 | 45.39 | 0.034 | 0.7107 | 1.0189 | 1.1402 |

**Location B**

| Element | Weight % | Atomic % | Error % | Net Int. | K Ratio | Z | A | F |
|---|---|---|---|---|---|---|---|---|
| C K | 74.09 | 81.82 | 5.73 | 819.74 | 0.4457 | 1.0216 | 0.5888 | 1 |
| O K | 20.33 | 16.86 | 12.69 | 116.24 | 0.0269 | 0.9764 | 0.1356 | 1 |
| Cl K | 0.97 | 0.36 | 10.86 | 47.14 | 0.0082 | 0.825 | 1.0006 | 1.0165 |
| Cu K | 4.6 | 0.96 | 8.87 | 49.01 | 0.0377 | 0.7123 | 1.0196 | 1.1277 |

**Location C**

| Element | Weight % | Atomic % | Error % | Net Int. | K Ratio | Z | A | F |
|---|---|---|---|---|---|---|---|---|
| C K | 76.5 | 83.01 | 5.12 | 1167.25 | 0.4964 | 1.017 | 0.6381 | 1 |
| O K | 19.75 | 16.09 | 12.84 | 140.18 | 0.0254 | 0.9718 | 0.1323 | 1 |
| Cl K | 0.79 | 0.29 | 9.85 | 49.23 | 0.0067 | 0.8206 | 1.0091 | 1.018 |
| Cu K | 2.97 | 0.61 | 11.96 | 42.06 | 0.0253 | 0.7082 | 1.0204 | 1.1803 |

**Figure S2.** EDS measurements at different locations on the copper grid containing Cu@HCl-$NH_2$ NPs. Images show only the carbon portions of the TEM grid (not the copper portions). The detection of copper on the carbon coating of the grids indicates that the nanoplatelets indeed contain copper

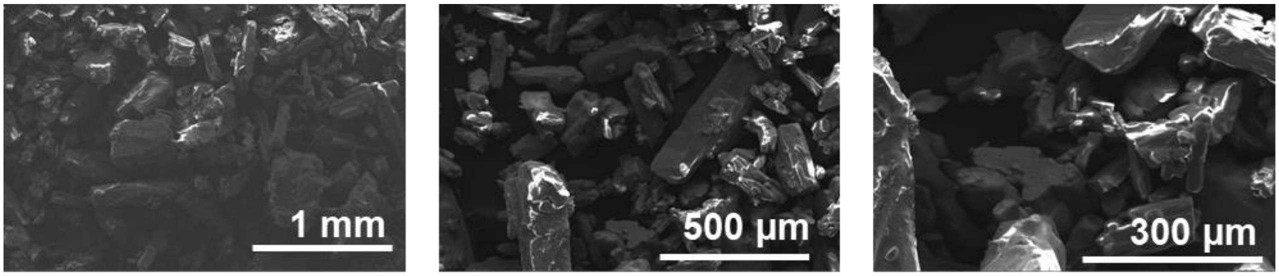


**Figure S3.** SEM images of commercially-obtained Cu(II) chloride powder.

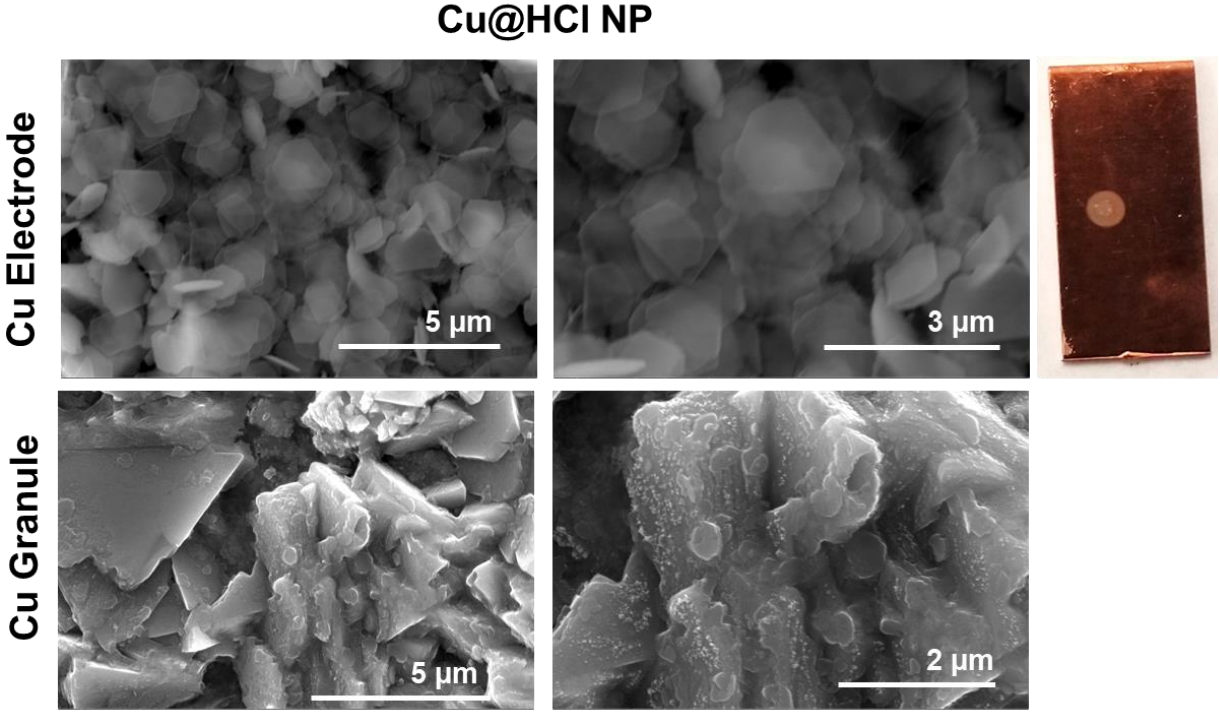


**Figure S4.** SEM images of Cu@HCl NPs assembled on a copper electrode and copper granule surface.

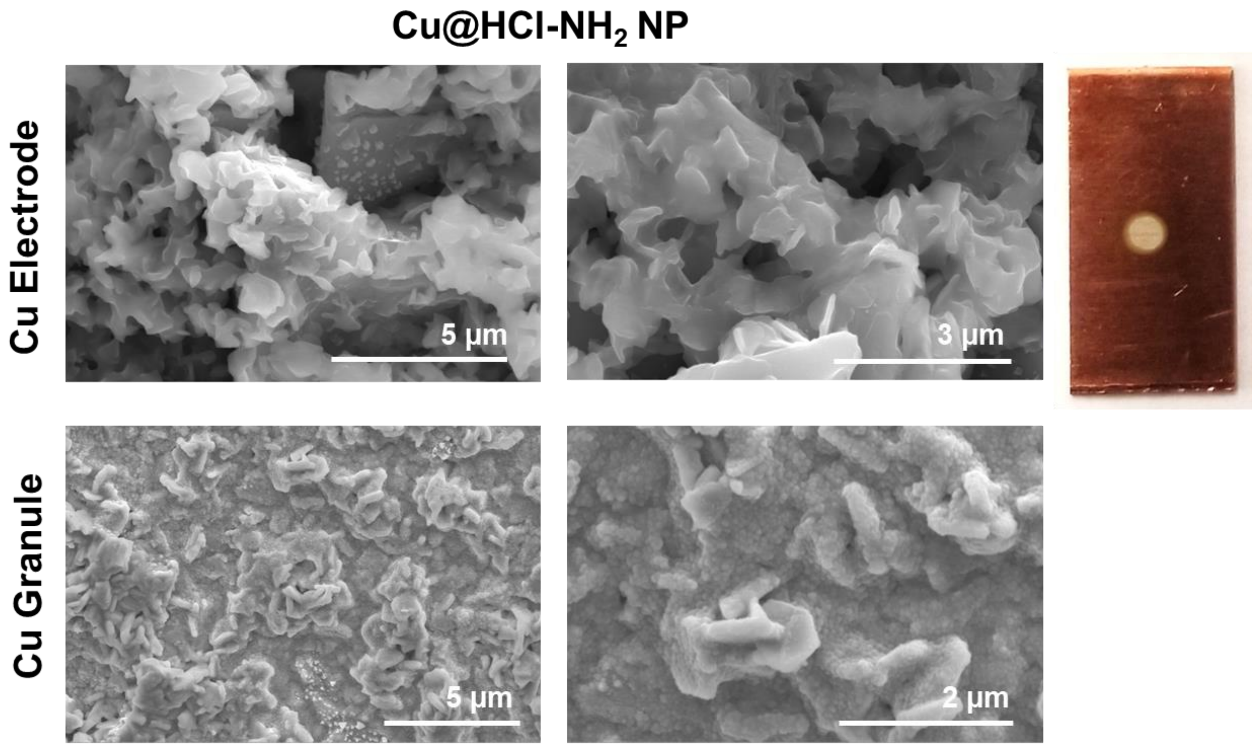


**Figure S5.** SEM images of Cu@HCl-NH$_2$ NPs assembled on a copper electrode and copper granule surface.

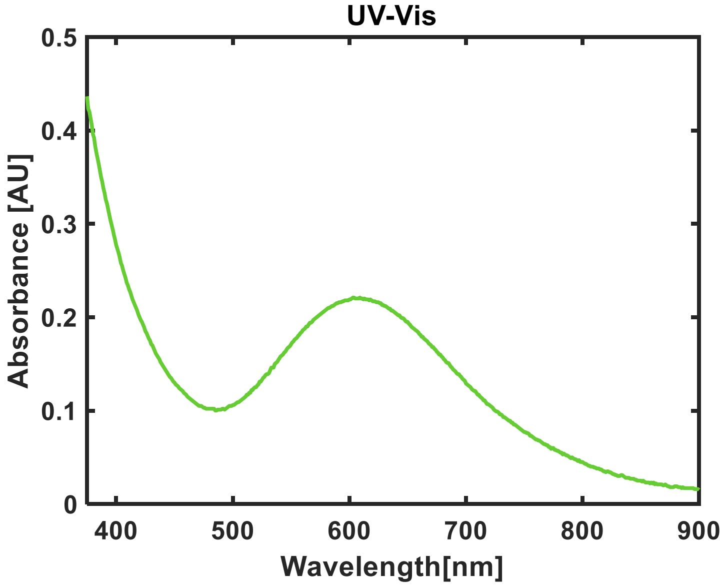


**Figure S6.** UV-visible absorbance spectrum of Cu@HCl-NH$_2$ NP dissolved in broth.

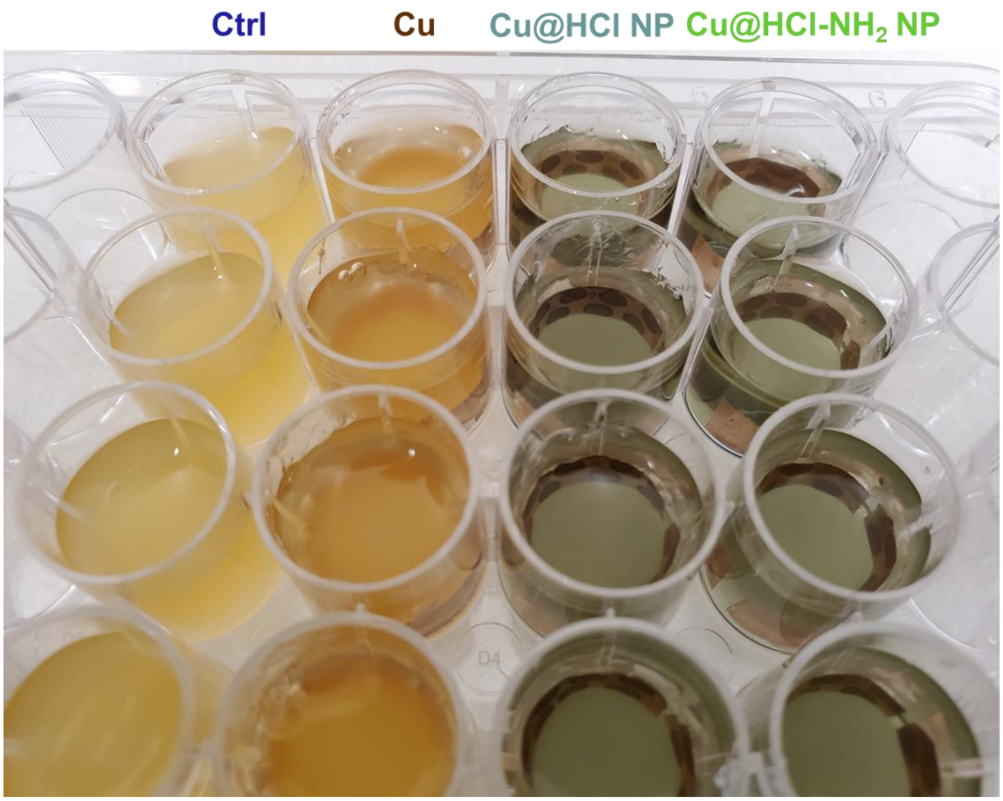


**Figure S7.** Photograph of bacterial wellplates after 4 h of treatment. A greater opacity is observed for the control and Cu-treated bacterial suspensions than the nanoplatelet-treated suspensions.

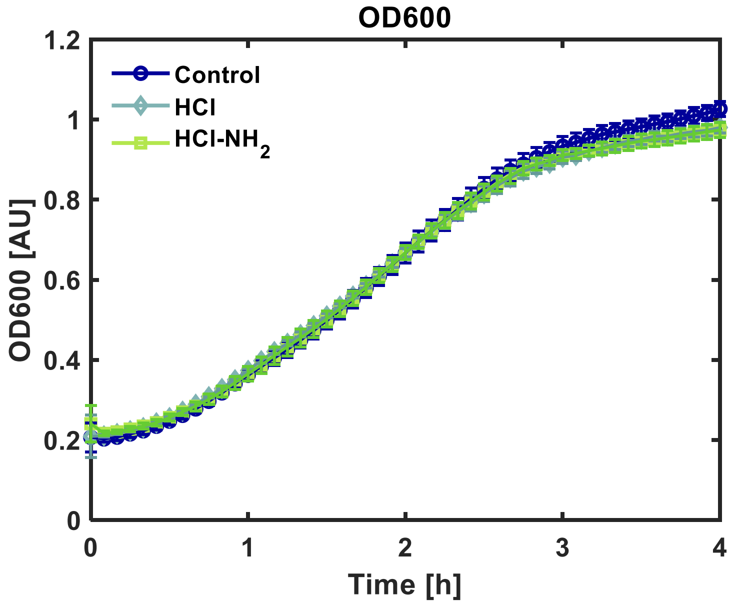


**Figure S8.** OD600 values for MDR E. Coli control (untreated) samples, as well as those treated with diluted HCl and HCl-$NH_2$ solutions. The solutions themselves appear to have little effect on bacterial cell division, suggesting that the nanoplatelets themselves, and not the synthesis solutions, are the cause of inhibited bacterial cell division. Values represent averages and error bars represent standard deviations (n=4).

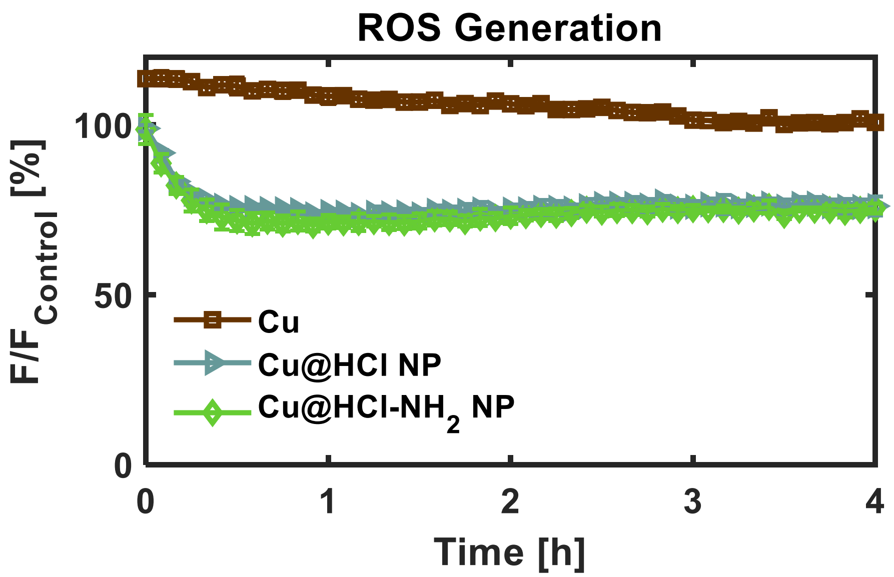


**Figure S9.** ROS Generation measured with time by fluorescence intensity of MDR E. Coli pre-incubated with CM-$H_2$DCFDA. Data are presented as a percent fluorescence intensity with respect to the measured intensity of the untreated control samples. Values represent averages and error bars represent standard deviations (n=4). A decrease in fluorescence intensity with respect to the control indicates there is not an increase in ROS generation.

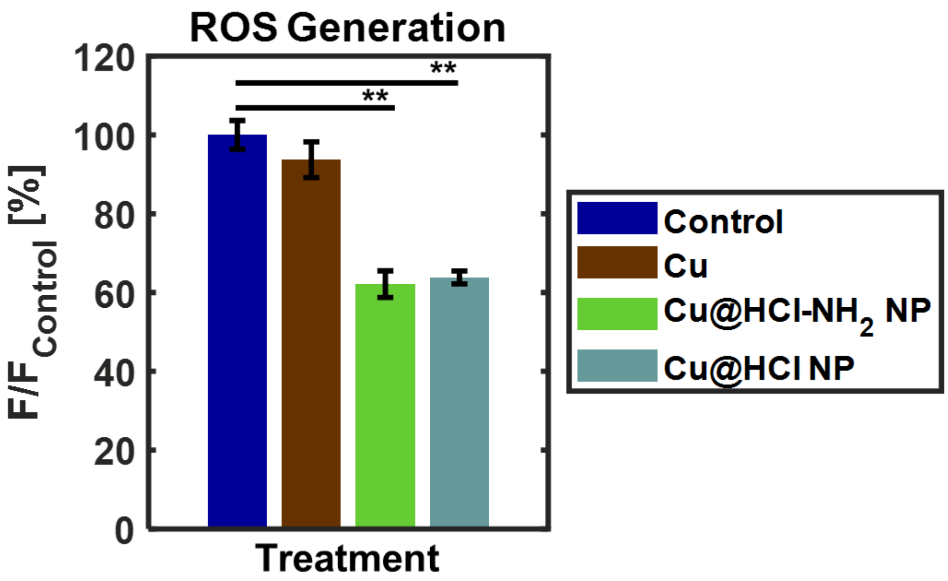


**Figure S10.** ROS Generation at four hours of treatment measured by fluorescence intensity of MDR E. Coli pre-incubated with CM-$H_2$DCFDA. Samples were protected from light for the 4 h treatment, after which fluorescence data was collected. Data are presented as a percent fluorescence intensity with respect to the measured intensity of the untreated control samples. Values represent averages and error bars represent standard deviations (n=4). ** indicates statistical significance with P<0.05. A decrease in fluorescence intensity with respect to the control indicates there is not an increase in ROS generation. These results confirm that the lower ROS generation observed in nanoplatelet-treated samples is not merely a result of photobleaching with time.

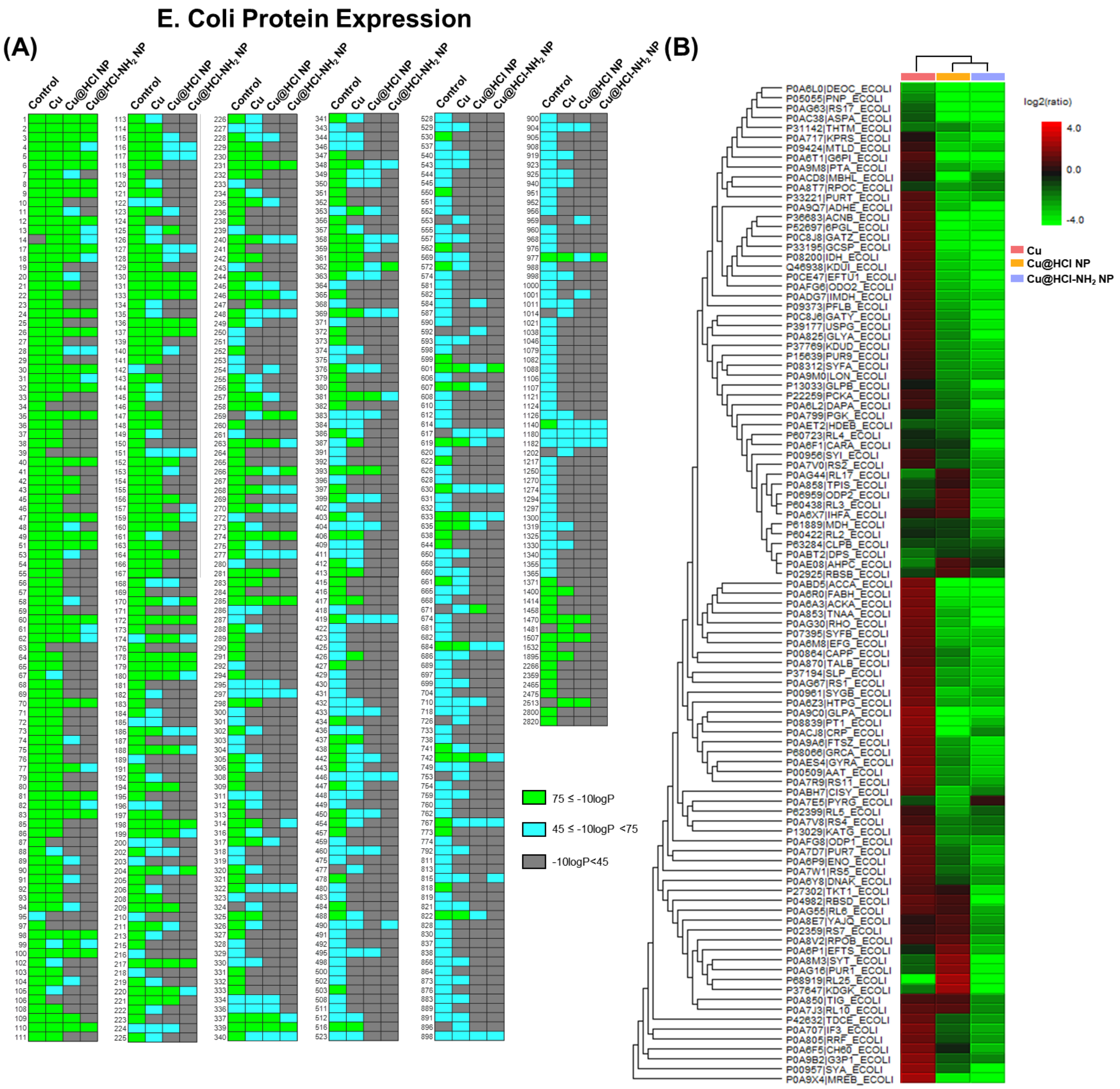


**Figure S11.** Proteomics analysis of E. Coli. **(A)** Heat maps classifying the -10logP values of each protein in each sample. Data represent a compilation of three technical replicates taken from each sample. **(B)** Hierarchical clustering of ratios between protein relative expression in each sample and protein relative expression in the control sample. Data represent the most data-rich technical replicate of three technical replicates for each sample.

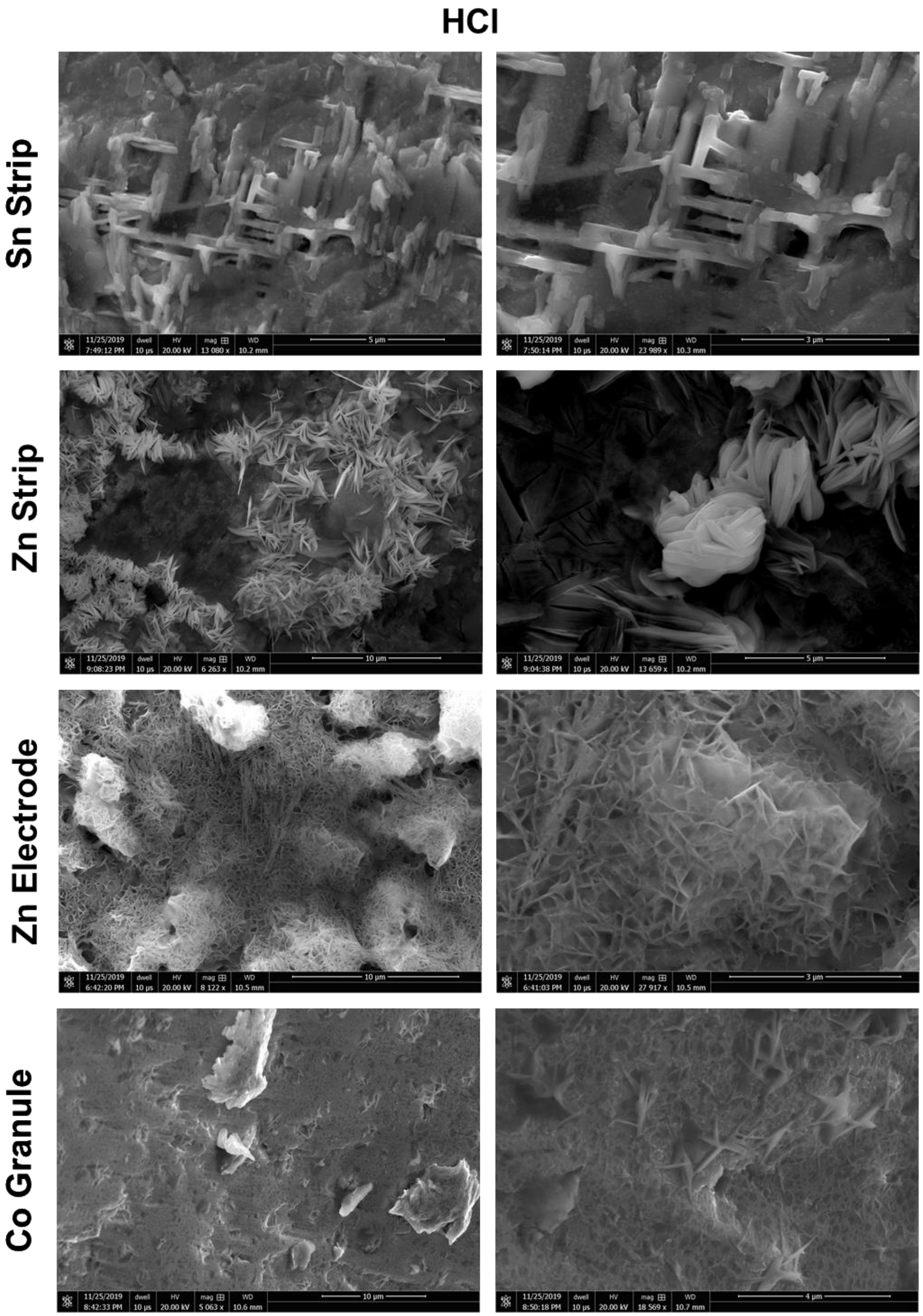


**Figure S12.** Nanostructures assembled using the HCl solution on Sn, Zn, and Co substrates.

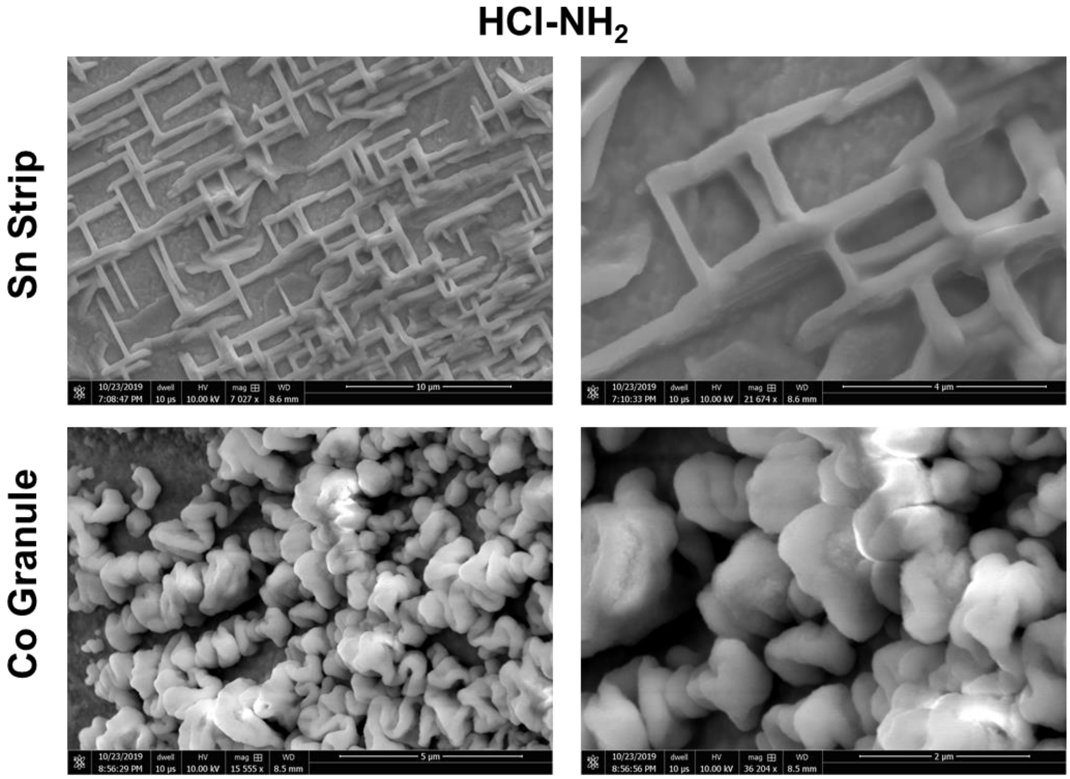


**Figure S13.** Nanostructures assembled using the HCl-$NH_2$ solution on Sn and Co substrates.
